# Placental mammal derived microRNAs alter pathways in the endometrial epithelia important for endometrial function

**DOI:** 10.1101/2022.04.23.489270

**Authors:** Laura Hume, Jessica C. Edge, Haidee Tinning, Dapeng Wang, Alysha S. Taylor, Vladimir Ovchinnikov, Annika Geijer-Simpson, Pavle Vrljicak, Jan J. Brosens, Emma S. Lucas, Nigel A.B. Simson, Jane Shillito, Karen Forbes, Mary J. O’Connell, Niamh Forde

**Author notes:** Both authors contributed equally to this work and can interchange their position when citing this paper (Laura Hume, Jessica C. Edge).

## Abstract

We tested the hypothesis that a panel of placental mammal-specific miRNAs and their targets play important to establish receptivity to implantation and their dysregulated expression may be a feature in women with early pregnancy loss. Relative expression levels of miR-340-5p, −542-3p, and −671-5p all increased following treatment of Ishikawa cells with progesterone (10 μg/ml) for 24 hrs (p < 0.05). RNA sequencing of these P4-treated cells identified co-ordinate changes to 6,367 transcripts of which 1713 were predicted targets of miR-340-5p, 670 of miR-542-3p, and 618 of miR-671-5p. Quantitative proteomic analysis of Ishikawa cells transfected with mimic or inhibitor (48 hrs: n=3 biological replicates) for each of the P4-regulated miRNAs was carried out to identify targets of these miRNAs. Excluding off target effects, mir-340-5p mimic altered 1,369 proteins while inhibition changed expression of 376 proteins (p < 0.05) of which, 72 were common to both treatments. A total of 280 proteins were identified between predicted (mirDB) and confirmed (*in vitro*) targets. In total, 171 proteins predicted to be targets by mirDB were altered *in vitro* by treatment with miR-340-5p mimic or inhibitor and were also altered by treatment of endometrial epithelial cells with P4. In vitro targets of miR-542-3p identified 1,378 proteins altered by mimic while inhibition altered 975 a core of 200 proteins were changed by both. 100 protein targets were predicted and only 46 proteins were P4 regulated. miR-671-mimic altered 1,252 proteins with inhibition changing 492 proteins of which 97 were common to both, 95 were miDB predicted targets and 46 were also P4-regulated. All miRNAs were detected in endometrial biopsies taken from patients during the luteal phase of their cycle, irrespective of prior or future pregnancy outcomes Expression of mir-340-5p showed an overall increase in patients who had previously suffered a miscarriage and had a subsequent miscarriage, as compared to those who had infertility or previous miscarriage and subsequently went on to have a life birth outcome. The regulation of these miRNAs and their protein targets regulate the function of transport and secretion, and adhesion of the endometrial epithelia required for successful implantation in humans. Dysfunction of these miRNAs (and therefore the targets they regulate) may contribute to endometrial-derived recurrent pregnancy loss in women.

## INTRODUCTION

Successful pregnancy in all placental mammals, including humans, is contingent on a carefully orchestrated series of events, including fertilisation between competent gametes, appropriate embryo development, successful implantation, and the establishment of a functional placenta. Perturbations at any of these key developmental points, such as chromosomal abnormalities in the embryo, are known to result in pregnancy loss. However, the contribution of an inadequately regulated endometrium is considerably underexplored. In humans, the uterine endometrium is a heterogeneous tissue composed of luminal and glandular epithelial cells, underlying fibroblast-like stromal cells, and populations of immune cells, vasculature and stem cells (1). Spatial and temporal responsiveness of these cells to hormonal cues and other signalling molecules throughout the menstrual cycle facilitates rapid proliferation and differentiation of the endometrial lining to prepare for successful implantation. The appropriate physiological response of the endometrium depends not only upon the cellular response to systemic cues, such as circulating oestradiol and progesterone (P4) levels, but also on the inter- and intra-cellular communication between different cell types.

Inappropriate remodelling of the endometrium, both prior to, and at the time of implantation, can lead to pregnancy failure such as miscarriage even when a developmentally competent embryo implants (2–4). Indeed, miscarriage can affect up to 1 in 6 pregnancies (5). The dynamic transcriptional responses of the endometrium during the menstrual cycle and implantation (reviewed by (6)) have been investigated in detail in relation to how they contribute to miscarriage (6–8). However, the regulatory signalling networks responsible for establishing a receptive endometrium to facilitate implantation, and their dysregulation in recurrent pregnancy loss (RPL), have yet to be determined.

Given their flexible and dynamic regulatory affects, miRNAs may be key in these processes. Once processed from pri- and pre-miRNA forms, these small RNA molecules are usually 22 nucleotides long and bind primarily to the 3’UTR of mRNAs and either suppress the initiation of translation or promote the degradation of the mRNA itself (9). They play key roles in inter-cellular communication as components of extracellular vesicles (EVs) or modify direct cargo loading to EVs (9). Some candidate microRNAs (miRNAs) that are altered in the endometrium have been reported to relate to early miscarriage (6–8). The expression of miRNAs varies dramatically and dynamically during pregnancy (10), with dysregulation of specific miRNAs also having been associated with other pregnancy disorders including preeclampsia and intrauterine growth restriction (11,12). However, conflicting data between studies, as well as cell-specific patterns of expression and regulation of miRNA mean we do not fully understand their role in normal and dysfunctional endometria.

The origin of placental mammals is synonymous with the emergence of placentation, lactation and also of hair. This burst of morphological innovation was mediated by a combination of novel and repurposed genomic elements (both protein coding and nonprotein coding), regulatory mechanisms, signalling pathways, cells and tissues (13–15). Indeed, bi-lateral signalling between foetal and maternal tissues is key to establishment of the mammalian placenta. In humans in particular, invasion of the maternal endometrium by the foetal trophoblast required co-ordinate specialisation of these distinct tissues, along with the co-evolution of the underlying molecular signalling networks facilitating these interactions and communications (13). Recent data using an evolutionary approach has identified a key role for the transcription factor, HAND2, and its target genes as important in the pathogenesis of pre-term birth and gestation length (16,17). We have adopted a similar strategy, leading to the identification of 13 miRNA families (17 microRNA genes) that arose coordinate with placental mammals and have been retained in all modern placental mammals (18). These miRNAs are detectable in the endometrial epithelium and their expression is altered in a species-specific manner by proteins known to facilitate signalling between the embryo and endometrium (19,20).

Given that 1) the endometrium and the trophoblast had to co-evolve regulatory networks to support implantation and early pregnancy success, 2) miRNAs can be important regulators in establishing these types of networks and 3) miRNA (and their targets) are regulated by molecules important to establish receptivity to implantation, we test the hypothesis that these mammal specific miRNA genes and their targets are regulated in the endometrium to facilitate receptivity to embryo implantation. Dysregulated expression of these miRNAs in the endometrium may be a feature in women with early pregnancy loss.

## MATERIALS AND METHODS

Unless otherwise stated all consumables were sourced from Sigma Aldrich (UK). The study was approved by the NHS National Research Ethics-Hammersmith and Queen Charlotte’s & Chelsea Research Ethics Committee (1997/5065).

### Cell culture

Human immortalised endometrial epithelial cells (Ishikawa cells - ECACC: #99040201) were cultured (50:50 DMEM/F12, 0.5 % GSP, 10 % charcoal-stripped FBS) at 37 °C and in 5 % CO_2_. Cells were cultured to 70 % confluence and seeded into 6-well plates. The cultures were treated with either 1) basal media (as above), 2) vehicle control (70 % ethanol, EtOH) or 3), 0.1 μg/ml P4, 4) 1.0 μg/ml P4, or 5) 1.0 μg/ml P4, for 24 hrs. Following treatment, cells were harvested by trypsinization and lysed in QIAzol Lysis Reagent. RNA was extracted using the miRNeasy Mini RNA Extraction kit (Qiagen, UK) as per manufacturer’s instructions.

### miRNA expression analysis and target prediction

Reverse transcription was carried out using the miRCURY LNA™1 RT Kit (Qiagen) using 10 ng of RNA according to manufacturer’s instructions, alongside reverse transcriptase negative controls. Expression analysis was performed using the miRCURY LNA™ SYBR Green PCR Kit 4000 (Qiagen) and a custom designed plate including miRNAs of interest and controls for normalisation (5S rRNA, U6 snRNA, UniSP3, and UniSP6). Thermal cycling and detection were performed on a ROCHE Light Cycler^®^ 480 II using the following parameters: 2 min at 90 °C, followed by 45 cycles of 10 secs at 95 °C & 60 secs at 56 °C, and melting curve analysis for 60 min at 95 °C. Cq values were determined and normalised to the geometric mean of normalisers. Data were analysed using ANOVA in Graphpad Prism to determine differences (determined as significant when p <0.05).

Predicted targets of miRNAs with P4-dependent expression were identified using TargetScan 7.2 (21), with output filtered based on the level of complementary binding between the miRNA seed region and target site: targets with 7mer or 8mer complementary binding to the seed region were selected, since the high number of consecutive matches is indicative of an efficient binding site (22). Ensembl transcript IDs provided by the TargetScan output were then mapped to gene names using Ensembl BioMart (release 98) (23).

### q-RT-PCR of predicted miRNA targets

Reverse transcription from 10 μg of RNA was carried out using the High-Capacity cDNA Reverse Transcription Kit (Applied Biosystems) as per the manufacturer’s protocol. Primers were designed for selected transcripts as described previously (20): Table 1) and 20 μM stock dilutions were prepared. PCR reactions were prepared in 20 μl volumes, containing 5 μl Roche SYBR Green Master Mix, 0.5 μl of each forward and reverse primer and 2 μl cDNA. Each sample was analysed in duplicate using the Lightcycler 480 II (Roche) as follows: 95 °C for 5 mins, 45 cycles of 95 °C for 10 secs, then 60 °C for 10 secs and finally 72 °C for secs, followed by melt curve analysis. Data were analysed using the comparative C_T_ method (2^-ΔΔct^) (24) normalised to the geometric mean of *ACTB, GAPDH* and *PPIA*. Significance differences were determined using ANOVA analysis in Graphpad Prism software, including Dunnett’s multiple comparisons test, to identify effects of treatment between each treatment concentration and the vehicle control. Statistical significance was assumed when p < 0.05 after multiple testing correction.

**Table 1.**
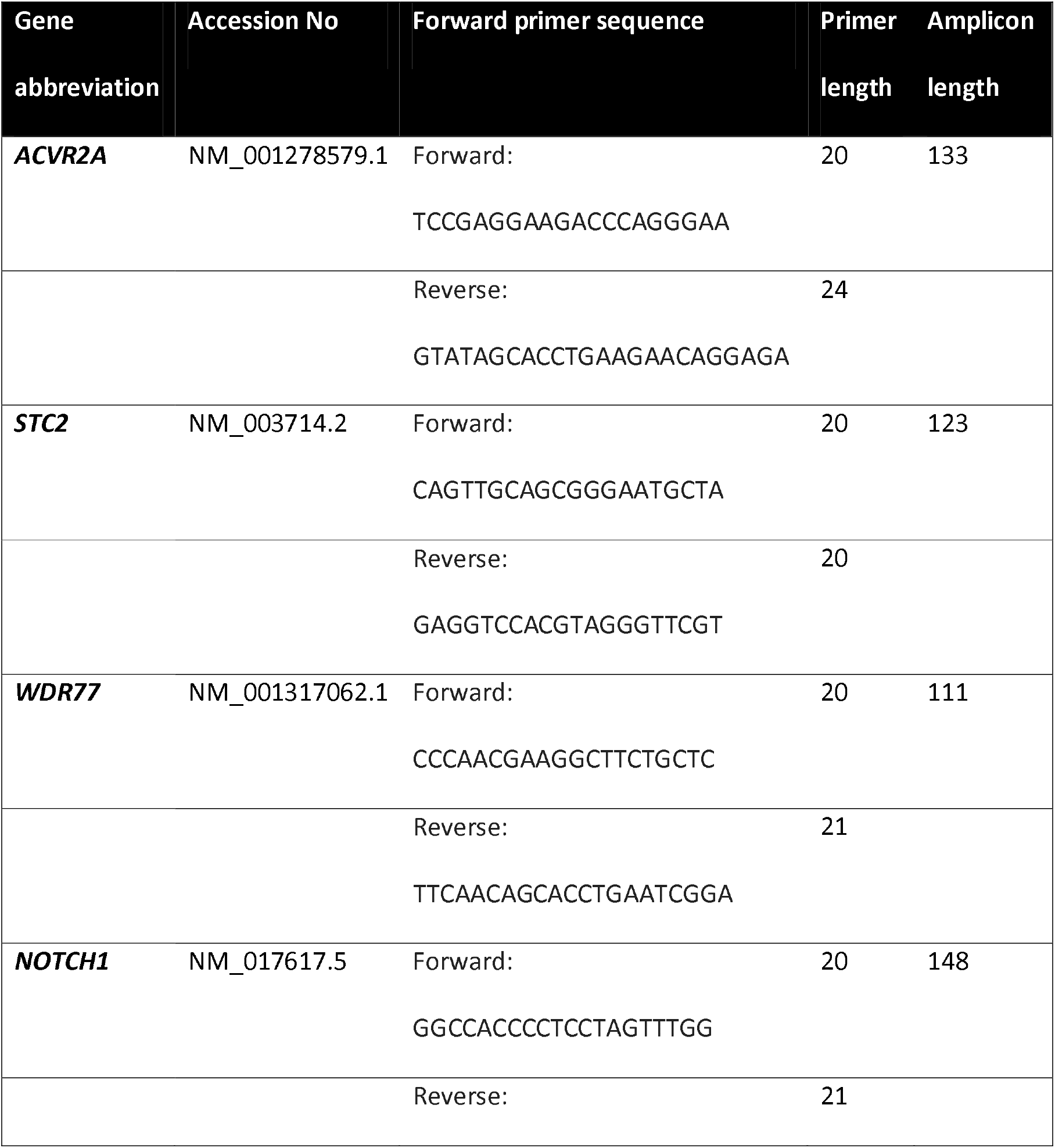
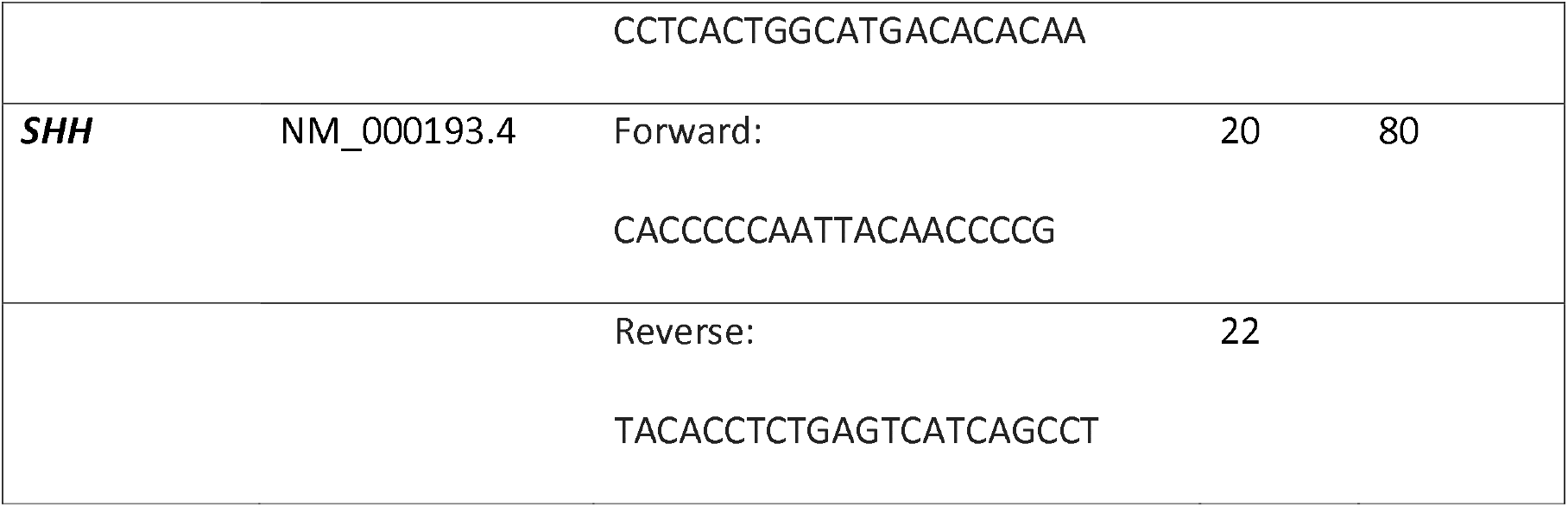
Primers used for q-RT-PCR analysis of P4 modified transcripts related to reproductive function. Gene name, accession number, forward primer sequence, length, reverse primer sequence, length, and amplicon length used for q-RT-PCR analysis of P4-regualted transcripts associated with reproduction function that are also predicted targets of P4-regualted miRNAs.

### RNA sequencing analysis

Total RNA (100 ng) was used for library preparation using the Illumina TruSeq^®^ Stranded Total kit, according to the manufacturer’s guidelines. The library quality, size range, and sequencing adaptor dimer contamination was assessed with an Agilent TapeStation 2200 using the DNA broad range kit. Excess sequencing adaptor dimer was removed by AxyPrep Mag PCR clean up Kit if present. Final libraries were quantified by the Qubit dsDNA assay kit and Qubit fluorometer (Life Technologies) before creating an equimolar pool of the libraries. The libraries were sequenced by the Next Generation Sequencing Facility at the University of Leeds, using the NextSeq 500 (Illumina, California, USA) to generate single end 75 bp reads. Reads were trimmed by Cutadapt (25) and filtered using fastq_quality_filter as part of FASTX-toolkit (http://hannonlab.cshl.edu/fastx_toolkit/) with parameters including “-q 20” and “-p 90”. The reference genome and gene annotation files of *Homo sapiens* were retrieved from GENCODE (release 31) (https://www.gencodegenes.org/) (26). Read mapping was performed by means of Subread aligners in Rsubread package (27) with only uniquely mapped reads reported in the final alignment and read summarisation. Principal Component Analysis (PCA) plot was carried out using all protein-coding genes and long noncoding RNAs with RPKM value ≥ 1 in at least one sample. Subsequently, log_2_(RPKM+1) transformation and quantile normalization were applied.

Quantification was carried out by featureCounts function in R. Significant differences in gene expression were identified using DESeq2 (28) with the following cut-offs: log_2_FoldChange > 1 (or < −1) and an FDR-adjusted P value of < 0.05. After this analysis, only protein-coding genes were retained based on the gene biotype labels. Overrepresentation enrichment analysis of DEGs were identified using WebGestalt (http://www.webgestalt.org/) (29) for gene ontology (GO) terms and pathways, with a significance enrichment level set at an FDR < 0.05.

### miRNA Mimic/inhibitor experiments

Ishikawa cells were plated (~200,000 cells/well) 24 hrs prior to treatment. At the time of treatment, cells were washed 3 times in PBS and 1.6 ml antibiotic & serum-free media added per well. The following treatments (n=3 biological replicates) were added 1) control medium, 2) Transfection reagent (Lipofectamine 2000), 3) miRNA mimic of interest (80 pmol/well), 4) miRNA inhibitor of interest, 5) miRNA non-targeting mimic, 6) miRNA non-targeting inhibitor. Cells were incubated for 48 hrs before media was removed, washed with PBS, and protein extracted using a RIPA buffer with protease inhibitors by passing the cells through a 21’G needle and syringe.

### Proteomic analysis

Fifty μg of protein from each sample, described above, was digested with trypsin (1.25 μg trypsin; 37 °C, overnight), labelled with Tandem Mass Tag (TMT) eleven plex reagents according to the manufacturer’s protocol (Thermo Fisher Scientific, Loughborough, LE11 5RG, UK) and the labelled samples pooled. One hundred μg of the sample pool was desalted using a SepPak cartridge according to the manufacturer’s instructions (Waters, Milford, Massachusetts, USA). Eluate from the SepPak cartridge was evaporated to dryness and resuspended in buffer A (20 mM ammonium hydroxide, pH 10) prior to fractionation by high pH reversed-phase chromatography using an Ultimate 3000 liquid chromatography system (Thermo Fisher Scientific). In brief, the sample was loaded onto an XBridge BEH C18 Column (130Å, 3.5 μm, 2.1 mm X 150 mm, Waters, UK) in buffer A and peptides eluted with an increasing gradient of buffer B (20 mM Ammonium Hydroxide in acetonitrile, pH 10) from 0-95 % over 60 mins. The resulting fractions (15 in total) were evaporated to dryness and resuspended in 1 % formic acid prior to analysis by nano-LC MSMS using an Orbitrap Fusion Lumos mass spectrometer (Thermo Scientific). In brief, peptides in 1 % (vol/vol) formic acid were injected onto an Acclaim PepMap C18 nano-trap column (Thermo Scientific). After washing with 0.5 % (vol/vol) acetonitrile, and 0.1 % (vol/vol) formic acid, peptides were resolved on a 250 mm × 75 μm Acclaim PepMap C18 reverse phase analytical column (Thermo Scientific) over a 150 mins organic gradient, using 7 gradient segments (1-6 % solvent B over 1 mins, 6-15 % B over 58 mins, 15-32 % B over 58 mins, 32-40 % B over 5 mins, 40-90 % B over 1 mins, held at 90 % B for 6 mins and then reduced to 1 % B over 1 mins) with a flow rate of 300 nl min^-1^. Solvent A was 0.1 % formic acid and Solvent B was aqueous 80 % acetonitrile in 0.1 % formic acid. Peptides were ionized by nano-electrospray ionization at 2.0 kV using a stainless-steel emitter with an internal diameter of 30 μm (Thermo Scientific) and a capillary temperature of 300 °C. All spectra were acquired using an Orbitrap Fusion Lumos mass spectrometer controlled by Xcalibur 3.0 software (Thermo Scientific) and operated in data-dependent acquisition mode using an SPS-MS3 workflow. FTMS1 spectra were collected at a resolution of 120 000, with an automatic gain control (AGC) target of 200 000 and a max injection time of 50 ms. Precursors were filtered with an intensity threshold of 5000, according to charge state (to include charge states 2-7) and with monoisotopic peak determination set to Peptide. Previously interrogated precursors were excluded using a dynamic window (60 secs +/-10 ppm). The MS2 precursors were isolated with a quadrupole isolation window of 0.7. m/z. ITMS2 spectra were collected with an AGC target of 10 000, max injection time of 70 ms and CID collision energy of 35 %. For FTMS3 analysis, the Orbitrap was operated at 50 000 resolution with an AGC target of 50 000 and a max injection time of 105 ms. Precursors were fragmented by high energy collision dissociation (HCD) at a normalised collision energy of 60 % to ensure maximal TMT reporter ion yield. Synchronous Precursor Selection (SPS) was enabled to include up to 10 MS2 fragment ions in the FTMS3 scan.

The raw data files were processed and quantified using Proteome Discoverer software v2.1 (Thermo Scientific) and searched against the UniProt Human database (downloaded January 2021: 169297 entries) using the SEQUEST HT algorithm. Peptide precursor mass tolerance was set at 10 ppm, and MS/MS tolerance was set at 0.6 Da. Search criteria included oxidation of methionine (+15.995 Da), acetylation of the protein N-terminus (+42.011 Da) and Methionine loss plus acetylation of the protein N-terminus (−89.03 Da) as variable modifications and carbamidomethylation of cysteine (+57.021 Da) and the addition of the TMT mass tag (+229.163 Da) to peptide N-termini and lysine as fixed modifications. Searches were performed with full tryptic digestion and a maximum of 2 missed cleavages were allowed. The reverse database search option was enabled and all data was filtered to satisfy false discovery rate (FDR) of 5 %. Protein groupings were determined by PD2.1, with minor modifications to allow inferrence of biological trends without any loss in the quality of identification or quantification. The MS data were searched against the human Uniprot database retrieved on 2021-01-14 and updated with additional annotation information on 2021-11-15. The protein abundances were normalised within each sample to total peptide amount, then Log2 transformed to bring them closer to a normal distribution. Statistical significance was then determined using paired *t*-tests between the conditions of interest with FDR corrected p-values obtained using the Benjamini-Hochberg method.

### Analysis of miRNAs in clinical samples

Human endometrial samples were obtained with written informed consent and in accordance with The Declaration of Helsinki (2000) guidelines. Biopsies were obtained from women attending the Implantation Clinic, a dedicated research clinic at University Hospitals Coventry and Warwickshire (UHCW) National Health Service Trust. Samples were timed relative to the pre-ovulatory LH surge, and obtained using a Wallach Endocell™ endometrial sampler. Following luteal phase endometrial biopsy, pieces of tissue (~2 mm) were preserved in RNA-later and stored at −80°C prior to analyses. Tissues were homogenised in 1 ml STAT-60 and RNA isolated according to the manufacturer’s instructions (AMS-Bio, UK). RNA was resuspended in 50 μl TE buffer (pH 8.0) and assessed by Nanodrop ND-1000 spectrophotometer, before storage at −80 °C. Samples underwent reverse transcription and miRNA expression analysis as described above. Expression of miRNAs of interest was measured in patients suffering from infertility and a subsequent live birth (IF+LB; n = 12), patients with miscarriage and a subsequent live birth (M+LB; n = 12), and patients suffering miscarriage and a subsequent miscarriage (M+M; n = 12).

## RESULTS

### Progesterone alters selected miRNAs and predicted targets in vitro

Relative expression levels of miR-340-5p, −542-3p, and −671-5p all increased following treatment with progesterone (10 μg/ml) for 24 hr (Figure 1) compared to control(s). No treatment effect was observed for the other miRNAs (p>0.05). TargetScan predicted miR-340-5p has binding sites in 6558 human transcripts. Of these predicted targets 68 % (4434) have functional assignments to ontologies/pathways in the interactome database, i.e. have an assigned function. For miR-542-3p, 4748 targets were predicted using TargetScan and 2041 have assigned functions in the reactome database. In the case of miR-671-5p 2820 of 4224 predicted targets have a function in the reactome database.

**Figure 1.**
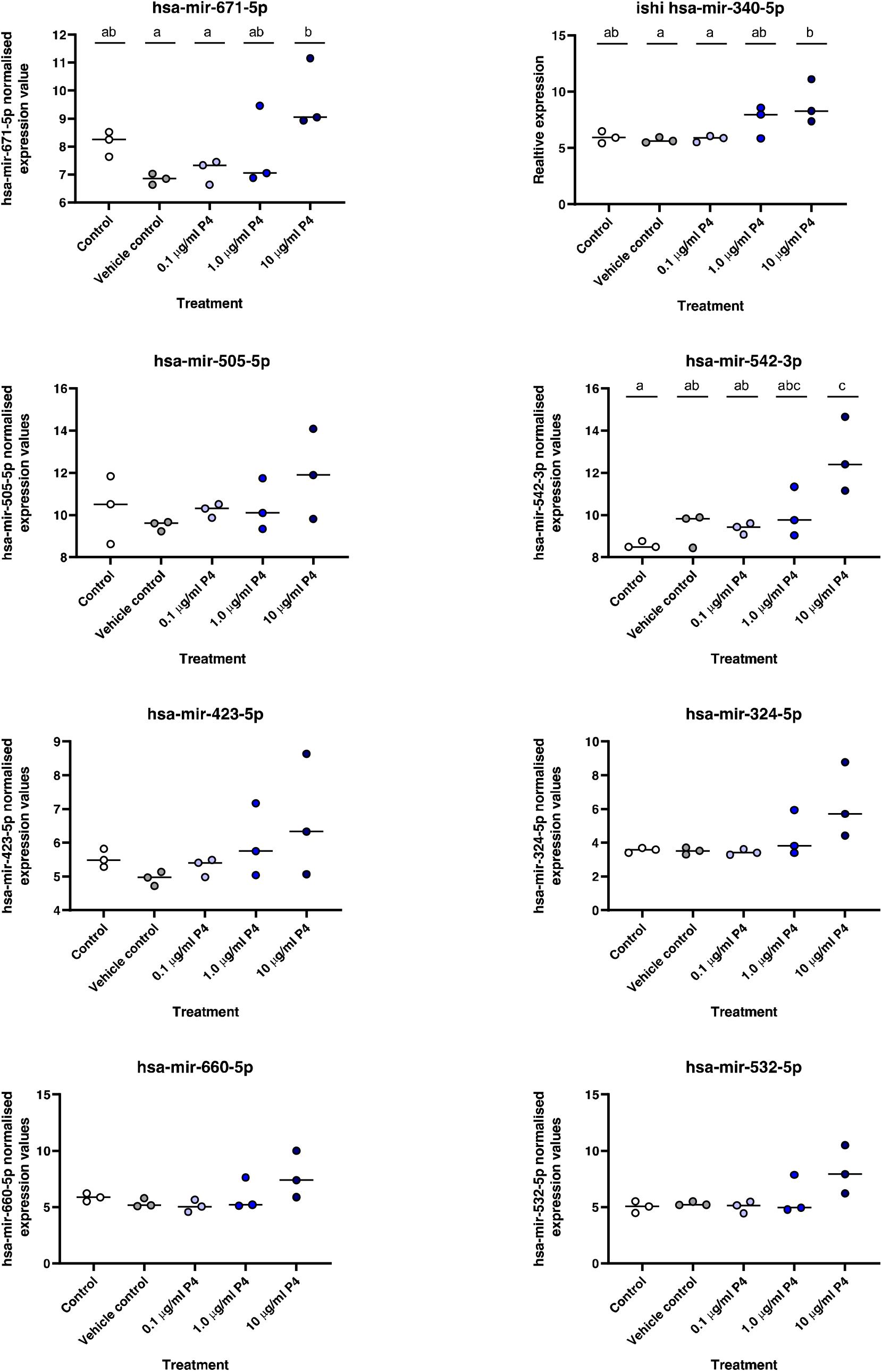
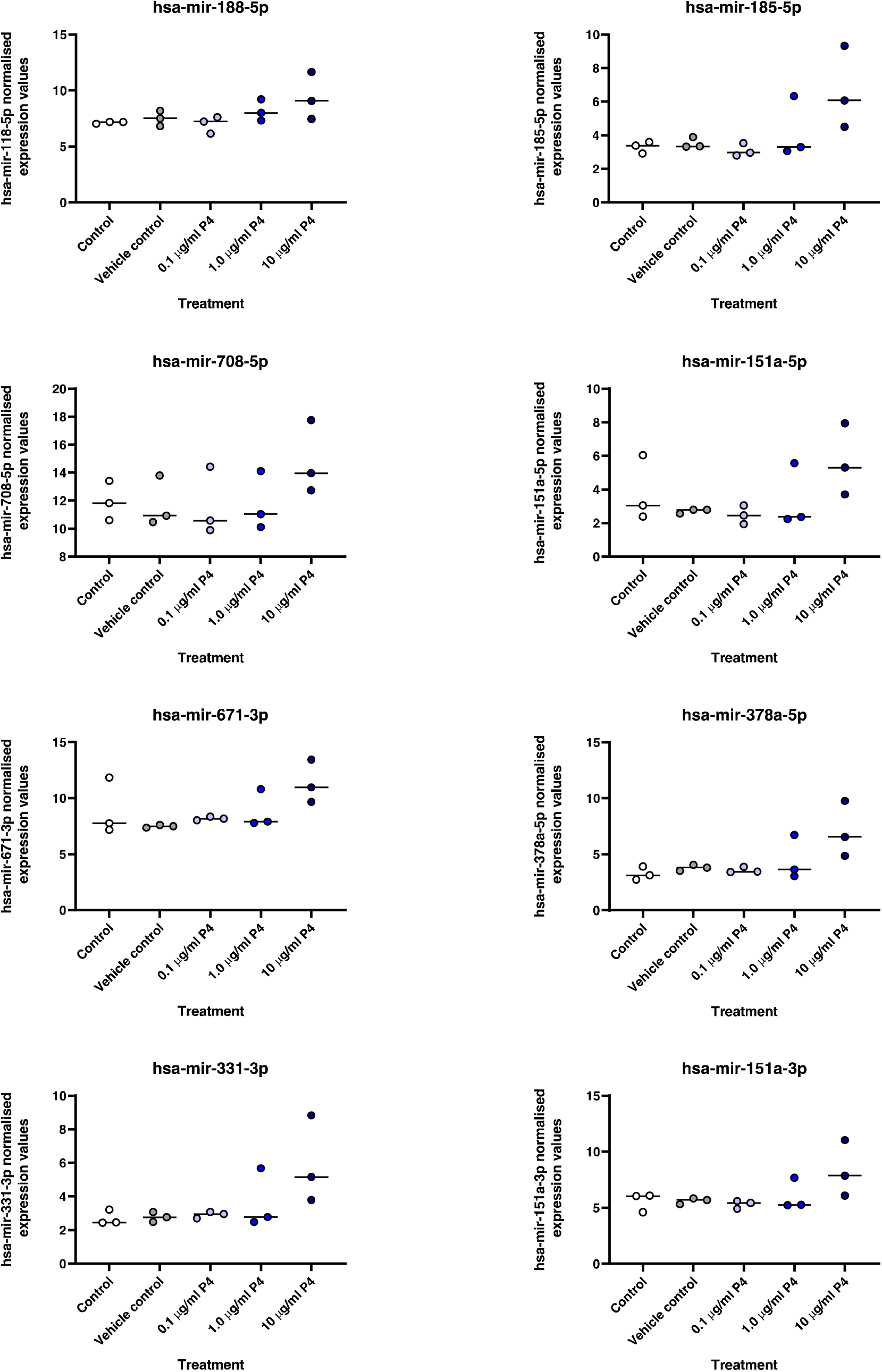
Expression of selected miRNAs in human endometrial epithelia (Ishikawa) cell line cultured for 24 hours with: 1) Control (open circle), 2) Vehicle Control (10% EtOH: grey circle), 3) 0.1ug/ml P4 (light blue circle), 4) 1.0 ug/ml P4 (royal blue circle), or 5) 10 ug/ml P4 (dark blue circle: n=3 biological replicates). Expression values of miRNAs were determined using LNA Sybrgreen method and the relative expression values following normalisation. Differences in expression were determined relative to <u>vehicle control</u> using an ANOVA and considered statistically different when P < 0.05. Differences between groups are denoted by difference in letters (a, b, c, d).

In total, 34 predicted targets of mir-340-5p, 20 predicted targets of mir-542-3p, and 16 predicted targets of mir-671-5p were associated with the GO term ‘reproductive function’ and were thus selected for further investigation. There was considerable overlap in the predicted ‘reproductive function’ targets across these three microRNA mature sequences where mir-542-3p and mir-671-5p had seven unique target predictions each (Figure 2A), conversely 19 were predicted to be targets of miR-340-5p alone. Common to all three P4-dependent miRNAs (i.e. mir-340-5p, mir-542-3p and mir-671-5p) were five ‘reproductive function’ predicted targets (Activin A Receptor, Type IIA [*ACVR2A*], Stanniocalcin-Related Protein [*STC2*], WD Repeat-Containing Protein 77 [*WDR77*], Notch Receptor 1 [*NOTCH1*], and Sonic Hedgehog Signalling Molecule [*SHH*], On examining these five common mRNA targets we found there was no effect of P4 on expression of *ARGHDIB, NOTCH1*, or *SHH* between any groups (Figure 2B: (p > 0.05)). Expression of *WDR77* increased in cells treated with the highest dose of P4 compared to controls (p < 0.05), while expression of *ACVR2A* expression decreased in cells treated with 1.0 and 10.0 ug/ml of P4 compared to control(s) (p < 0.05).

**Figure 2A.**
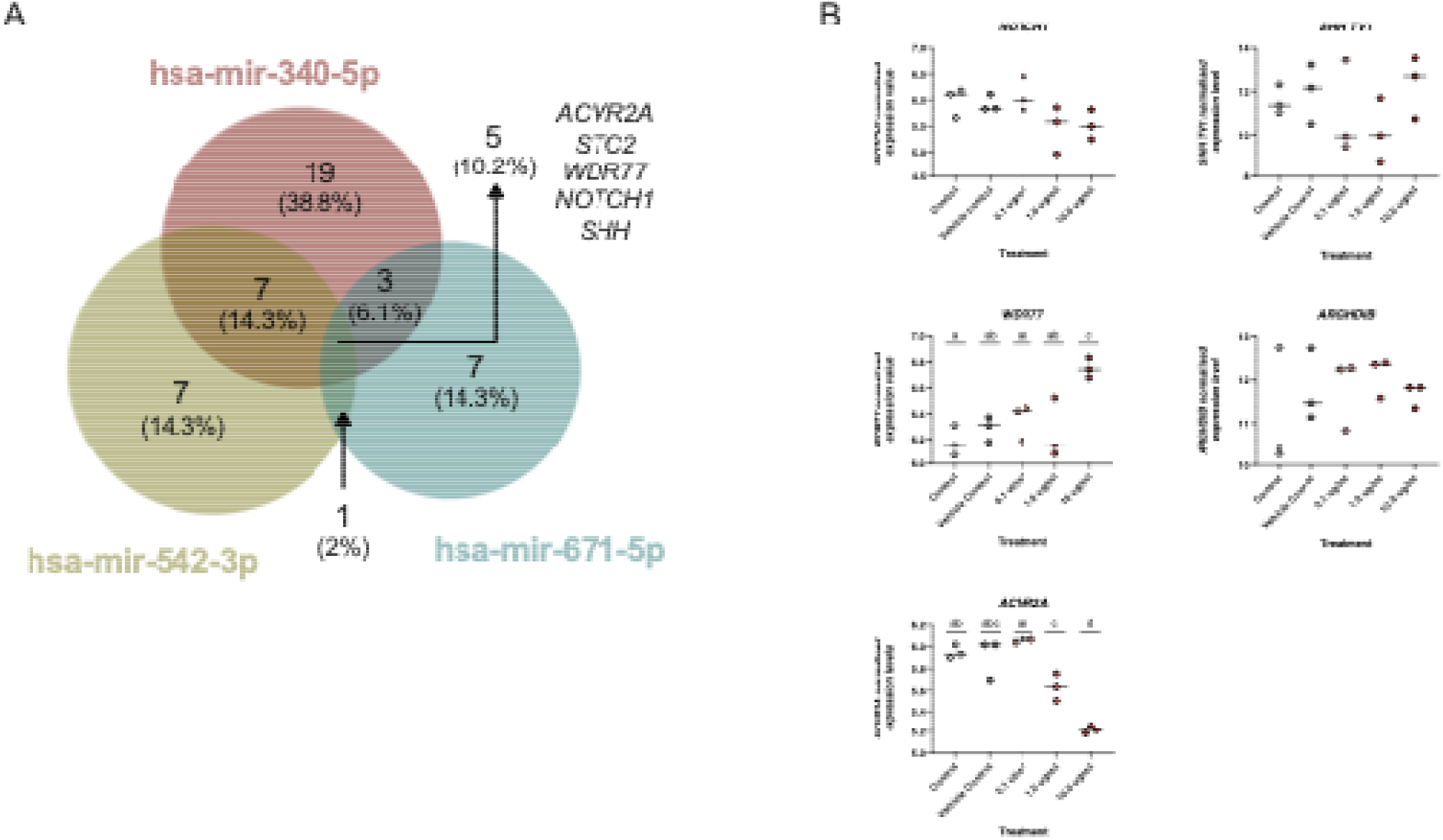
Venn diagram representing overlap in predicted mRNA targets of the three miRNAs altered following 24 hr treatment of endometrial epithelial cells with P4 *in vitro*. These were associated with the Gene Ontology Biological Process of Reproductive Biology. **2B.** RT-qPCR analysis of predicted targets of all three P4-regulated miRNAs that are associated with the Gene Ontology Biological Process of Reproductive Function. Expression was determined in Ishikawa cells treated with control (open circles), vehicle control (grey circles), 0.1 μg/ml (pale red circle), 1.0 μg/ml (medium red circle), or 10.0 μg/ml (dark red circle) of P4 for 24 hrs *in vitro* (n=3 independent cultures). Differences in expression compared were determined using ANOVA with differences depicted by different subscripts (a, b, c, d) when p < 0.05.

### Coordinate changes in mRNAs in endometrial epithelial cells treated with progesterone

Next, we sought to identify co-ordinate changes in mRNA expression occurring with the changes in miRNA expression at the highest concentration of P4. Principal Component Analysis (PCA) of RNA sequencing data revealed clear separation between controls and P4 treated cells (Figure 3A). Following data quality control step and a Benjamini-Hochberg FDR correction, differentially expressed protein coding genes (DEGs) were determined. In total, 6,367 DEGs were identified, 1,238 of which had a log2 fold change of greater than 1 (Supplementary Table 1).

**Figure 3A.**
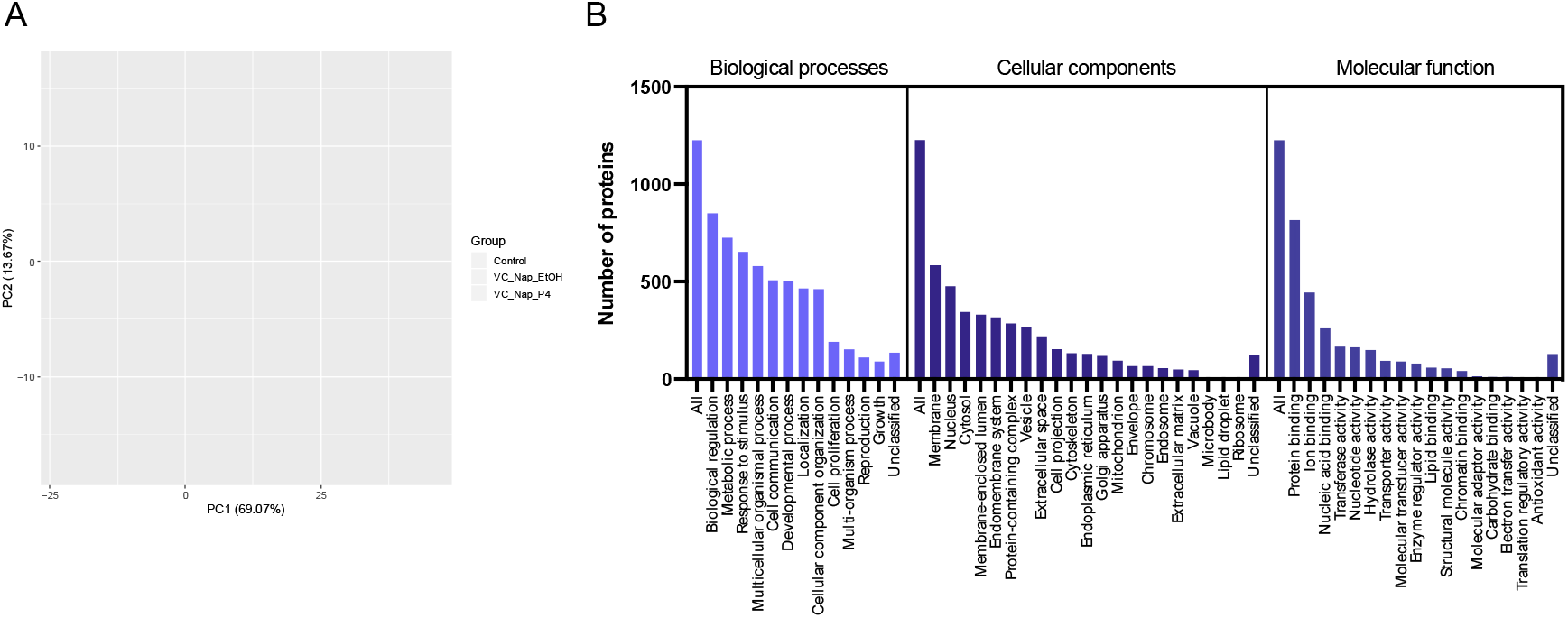
RNA sequencing analysis of Ishikawa cells treated with P4 in vitro. Principal component analysis depicting the overall transcriptional differences between cells via RNA sequencing of Ishikawa cells treated with 1) Control (orange circle), 2) Vehicle control (green circle), and 3) 10 ug/ml P4 (blue circle) for 24 hours. Controls clearly separate on the left-hand side while P4 treatment clearly cluster together on the right-hand side of the graph (n=3 biological replicates per group). B. Bar chart of Biological Processes (red), Cellular Component (blue), and Molecular Function (green) associated with P4 modified transcripts.

P4 altered the expression of transcripts in the categories of: biological regulation, metabolic processes, localised to the membrane and to the nucleus, and molecular functions of protein, ion, and nucleic acid binding more than was expected by chance (Figure 3B). Biological processes enriched in these P4 treated cells included Notch signalling pathway, tube development, tube morphogenesis, and epithelium development (Figure 4A: Supplementary Table 2). Cellular components were enriched in ‘cell-substrate junction’, and ‘integral component of plasma membrane’ (Figure 4B: Supplementary Table 3), while molecular functions influenced by P4 treatment included DNA-binding transcription activator activity, RNA polymerase II regulatory region sequence-specific interactions (Figure 4C: Supplementary Table 4). The signalling pathways that were enriched involved signalling by NOTCH1, amino acid transport and DNA methylation (Figure 5A: Supplementary Table 5) while enriched KEGG pathways were involved in p53 signalling, Ferroptosis, Melanoma, and Glioma (Figure 5B: Supplementary Table 6). The gene ontology enrichment network used a total of 22,194 genes with 1,089 seeds in the selected network. A total of 844 seeds were identified in the expanded sub-network and they produced an enriched biological process network including those involved in developmental processes, epithelial development, and both positive and negative regulation of cellular processes (Figure 5C).

**Figure 4.**
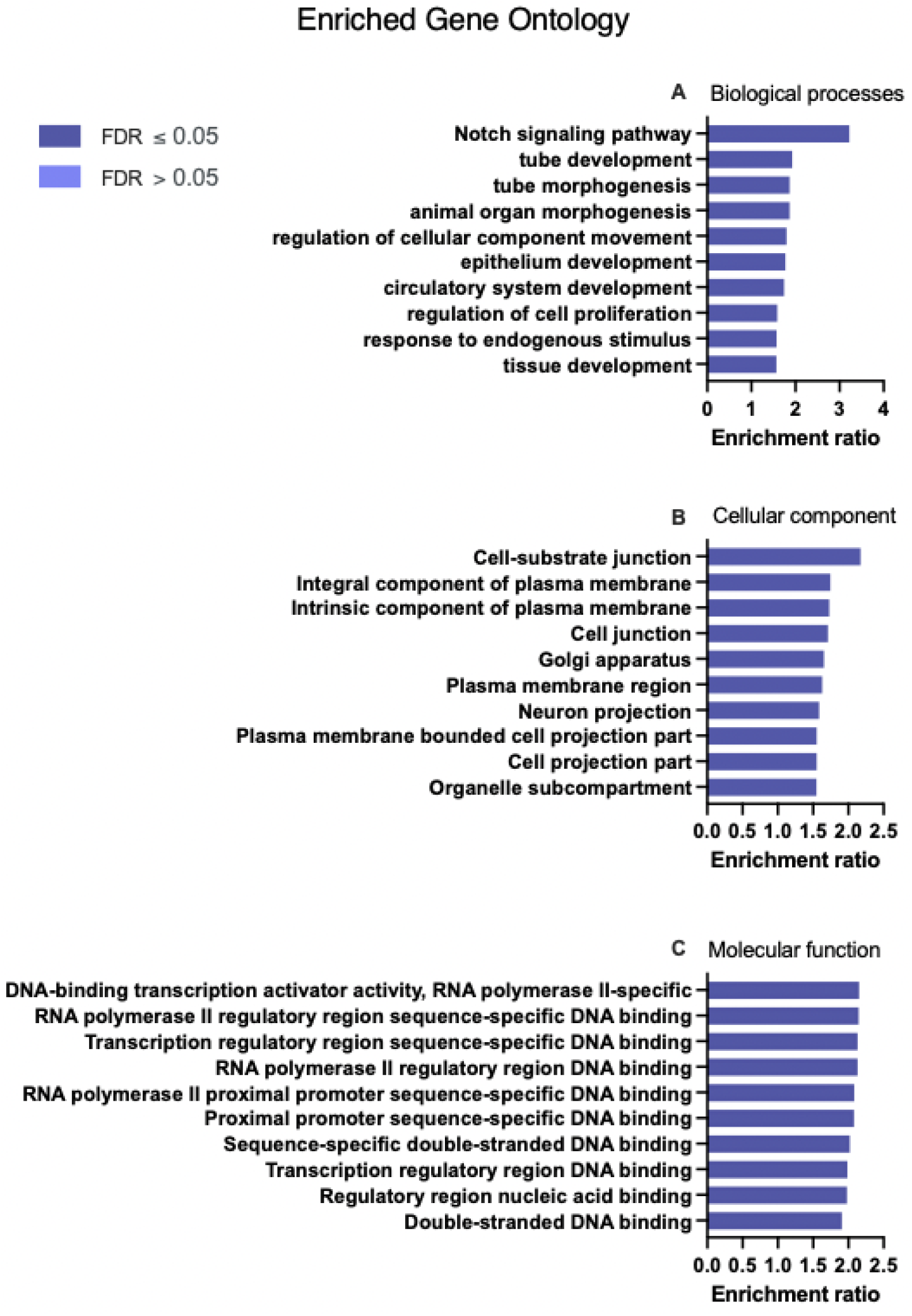
Enriched Gene Ontology **(A)** Biological Processes, **(B)** Cellular Component, and **(C)** Molecular Functions associated with P4 modified transcripts determined following RNA sequencing of human endometrial epithelial cells (Ishikawa cells) treated with 10 ug/ml P4 for 24 hours. These are significantly enriched i.e. more transcripts associated with these ontologies than one would expect by chance (FDR <0.05).

**Figure 5.**
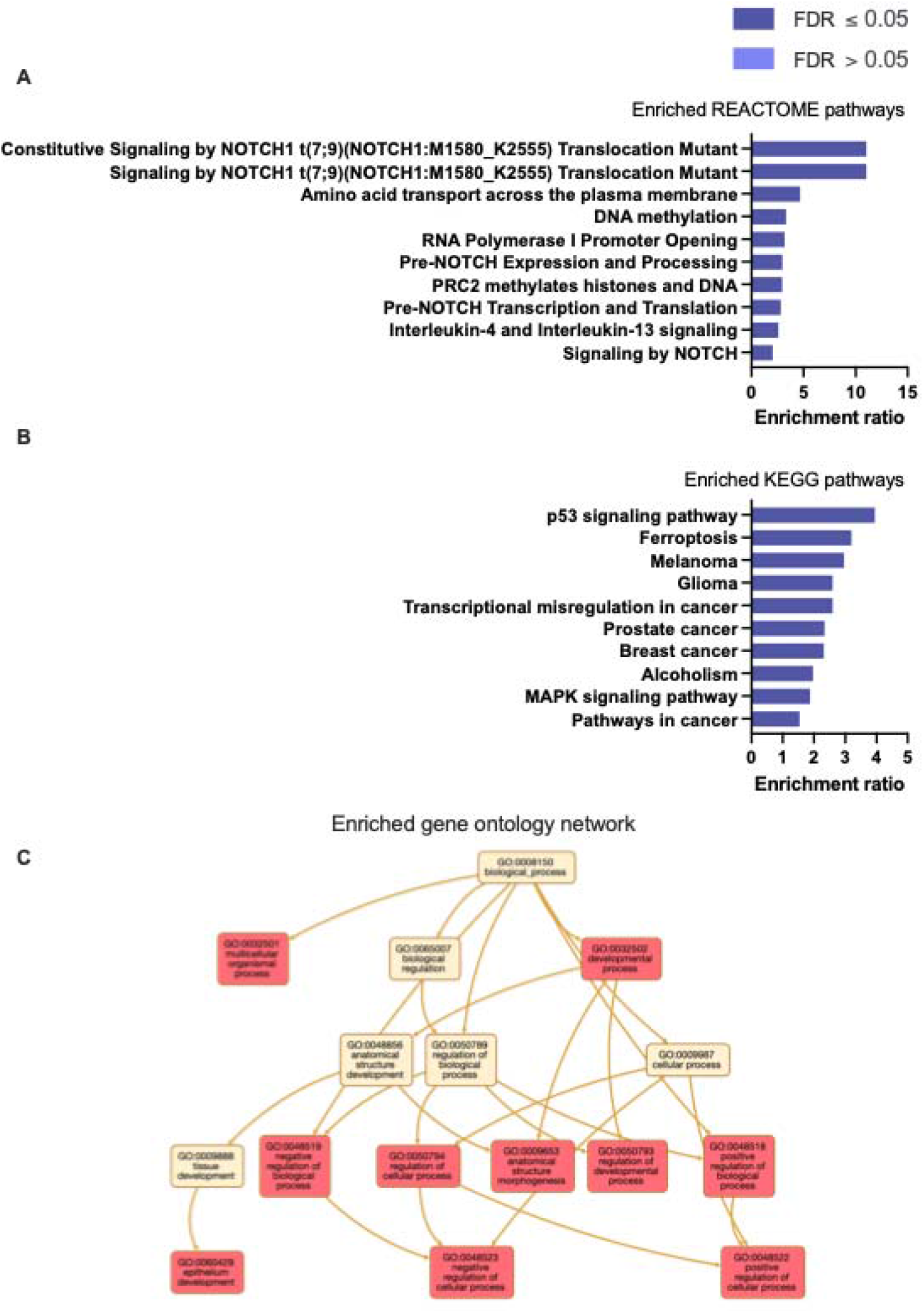
Enriched **(A)** REACTOME pathways, **(B)** KEGG pathways, and **(C)** ontology network, associated with P4 modified transcripts determined following RNA sequencing of human endometrial epithelial cells (Ishikawa cells) treated with 10 ug/ml P4 for 24 hours. These are significantly enriched i.e. more transcripts associated with these pathways than one would expect by chance (FDR <0.05).

**Figure 5.**
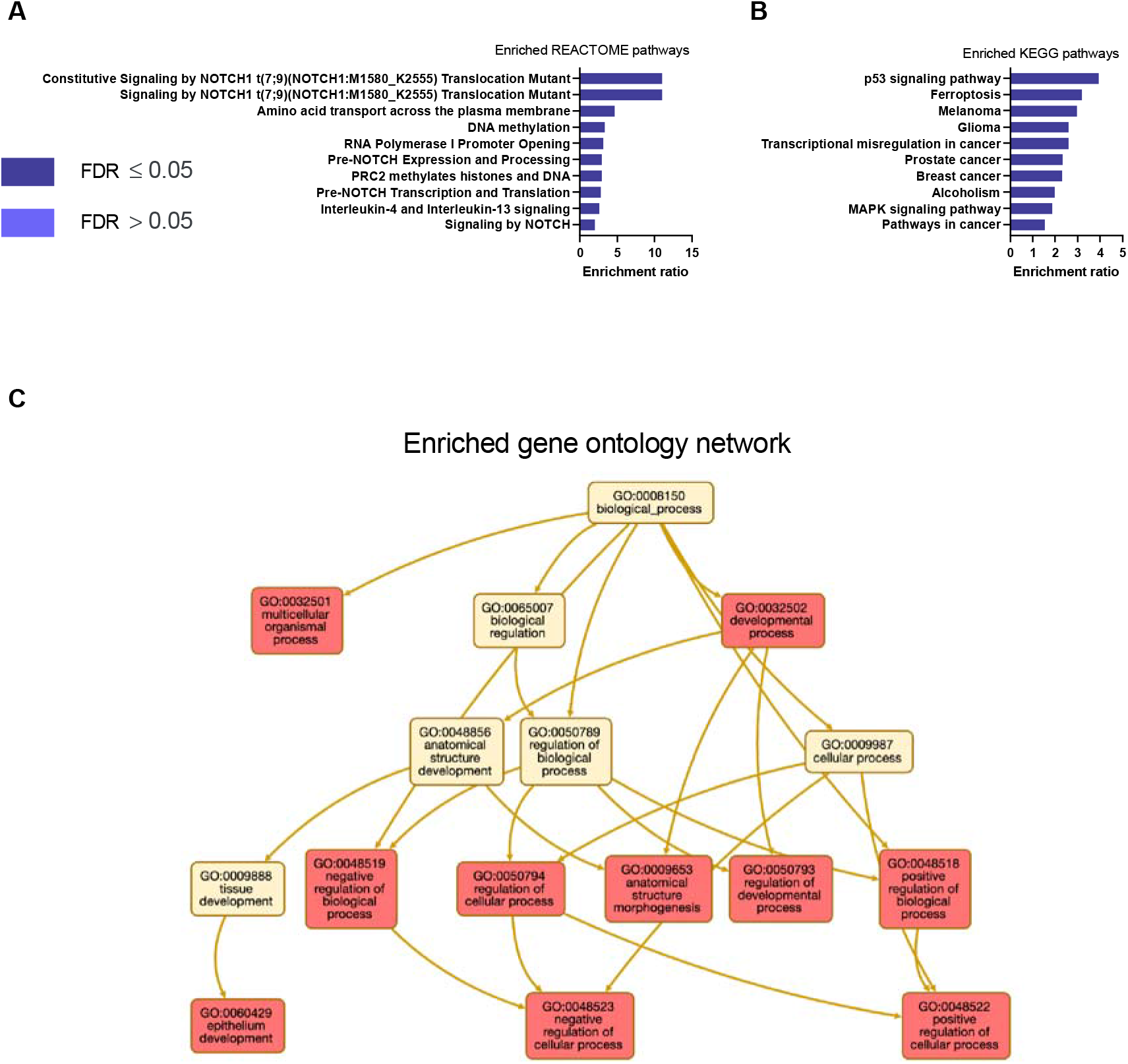
Enriched **(A)** REACTOME pathways, **(B)** KEGG pathways, and **(C)** ontology network, associated with P4 modified transcripts determined following RNA sequencing of human endometrial epithelial cells (Ishikawa cells) treated with 10 ug/ml P4 for 24 hours. These are significantly enriched i.e. more transcripts associated with these pathways than one would expect by chance (FDR <0.05).

We then investigated the overlap in the P4-dependent expression profiles and the predicted targets from Targetscan of the three P4-regulated miRNAs (Figure 6). Of the predicted targets of miR-340-5p: 1713 transcripts were also changed in P4 treated cells; miR-542-3p: 670 transcripts, and miR-671-5p: 618 transcripts indicating distinct P4-miRNA transcriptomic signatures. Complete details of transcripts are provided in Supplementary Table 7. A cohort of 473 transcripts that are predicted targets of all three P4-regulated miRNAs exhibit P4-responsive expression in the same cells. No significantly enriched biological processes or KEGG/panther pathways were identified (Figure 7A; Supplementary Table 8 and Figure 8; Supplementary Tables 11 & 12). However, significant enrichment for cellular components including those involved in the nuclear membrane, nuclear envelope, and endosome (Figure 7B: Supplementary Table 9) were observed. Molecular functions of ubiquitin-ubiquitin ligase activity, protein serine/threonine kinase activity, and adenyl nucleotide activity were also significantly enriched (Figure 7C: Supplementary Table 10).

**Figure 6.**
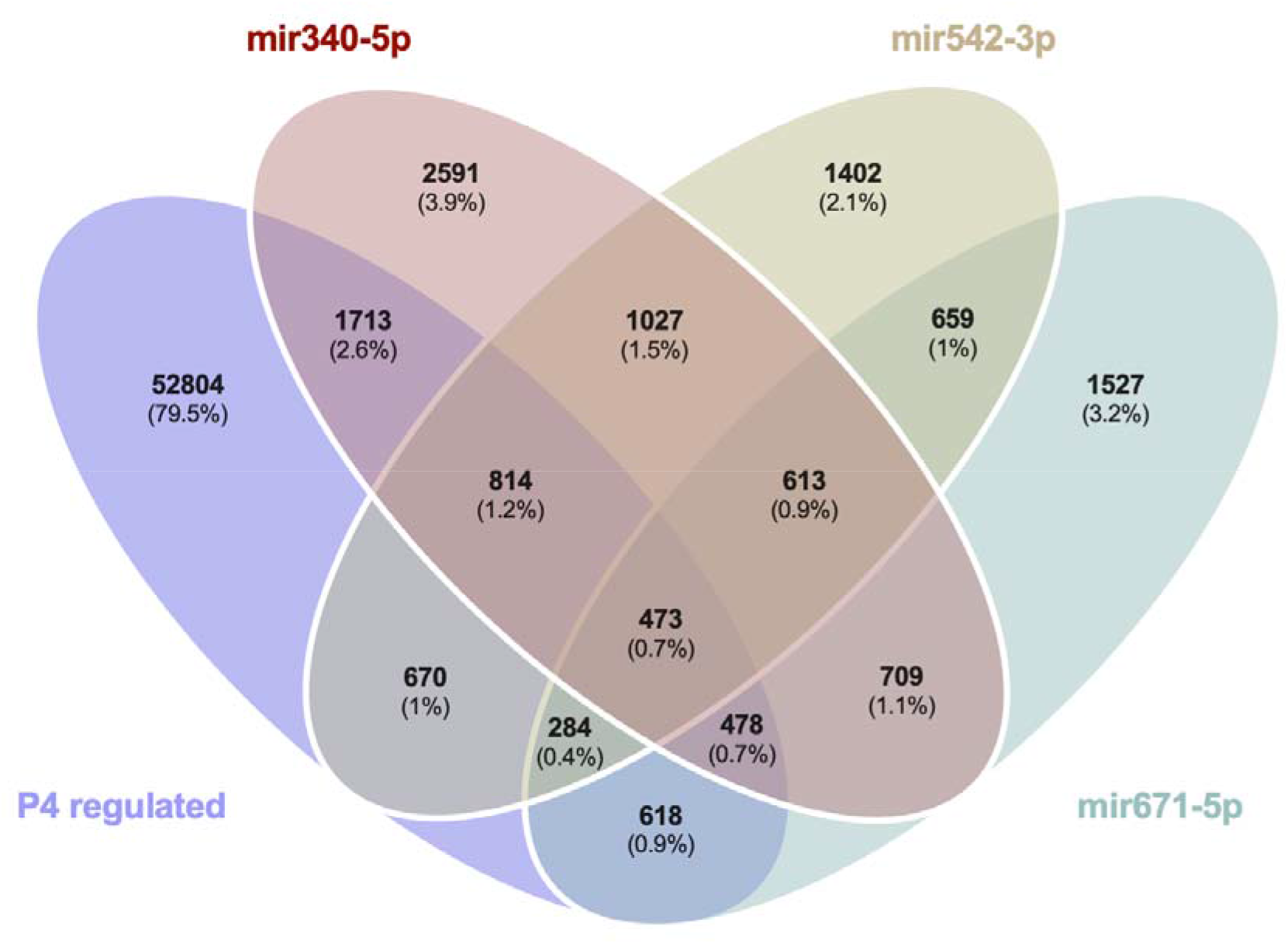
Venn diagram analysis demonstrating the overlap in predicted targets (from miRbase) of the 3 miRNAs modified by treatment of Ishikawa cells with P4 for 24 hrs in vitro as well as those mRNAs that were identified as altered in the same cells following P4 treatment for 24 hours compared to vehicle controls.

**Figure 7.**
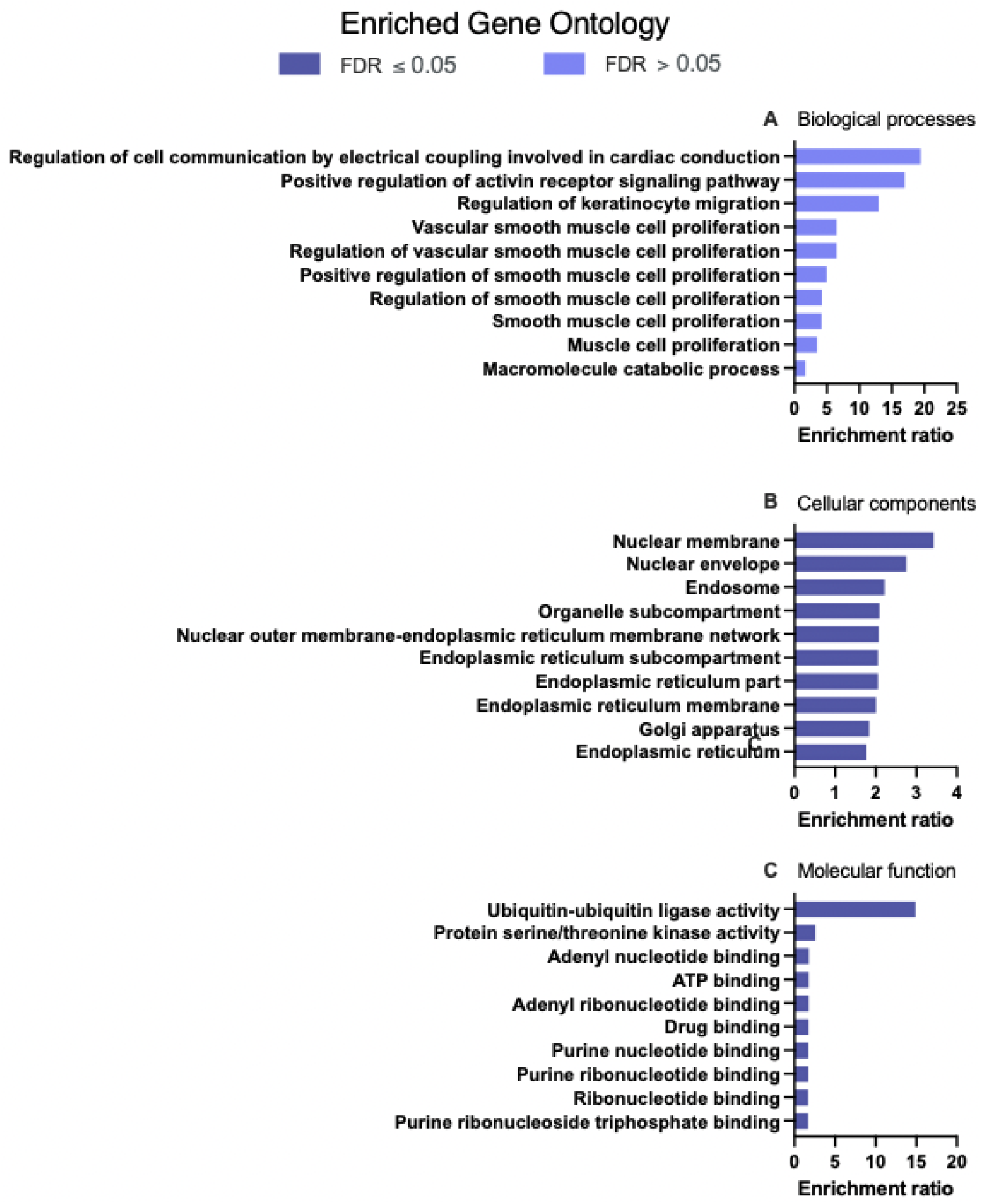
Enriched Gene Ontology **(A)** Biological Processes, **(B)** Cellular Component, and **(C)** Molecular Functions associated with 473 P4 modified transcripts that are also predicted targets of the 3 miRNAs modified in human endometrial epithelial cells (Ishikawa cells) treated with 10 ug/ml P4 for 24 hours. These are significantly enriched i.e. more transcripts associated with these ontologies than one would expect by chance (FDR <0.05).

**Figure 8.**
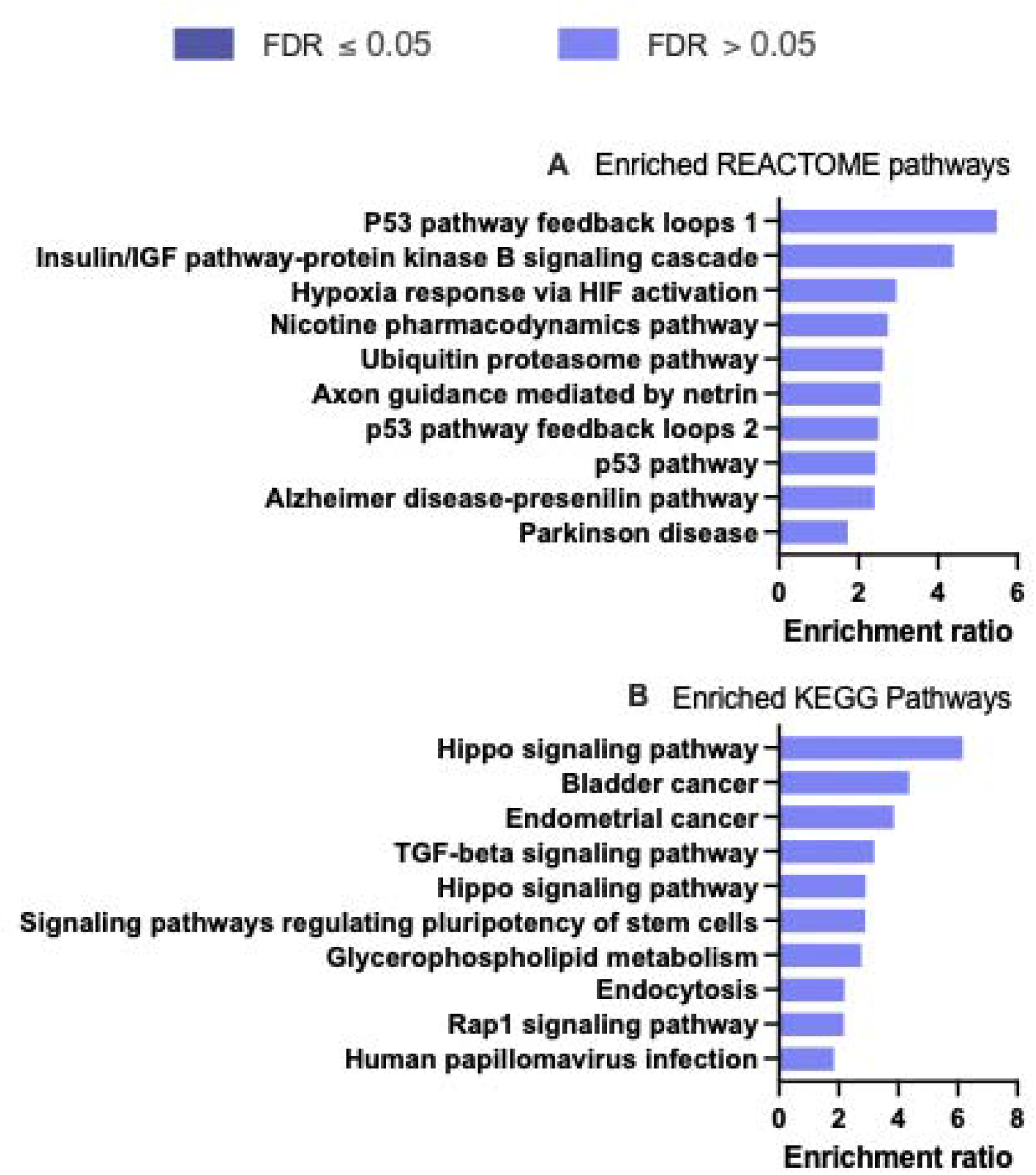
Enriched **(A)** REACTOME pathways, and **(B)** KEGG pathways associated with 473 P4 modified transcripts that are also predicted targets of the 3 miRNAs modified in human endometrial epithelial cells (Ishikawa cells) treated with 10 ug/ml P4 for 24 hours. These are significantly enriched i.e. more transcripts associated with these ontologies than one would expect by chance when FDR <0.05.

### In vitro targets of miR-340-5p in endometrial epithelial cells

Excluding off target effects, treatment of endometrial epithelial cells with mir-340-5p mimic altered 1,369 proteins as compared to controls (p < 0.05; Supplementary Figure 1; Supplementary Table 22). These were significantly enriched in pathways involved in oxidation-reduction function and metabolism pathways (Supplementary Figure 1). In contrast, inhibition of mir-340-5p changed expression of 376 proteins (p < 0.05; Supplementary Figure 2; Supplementary Table 23) including those involved in tRNA and ncRNA processing as well as nuclear transport (Supplementary Figure 2). A cohort of 72 proteins were altered in endometrial epithelial cells following treatment with both mimic and inhibitor of mir-340-5p (Figure 9; Supplementary Table 13).

**Figure 9:**
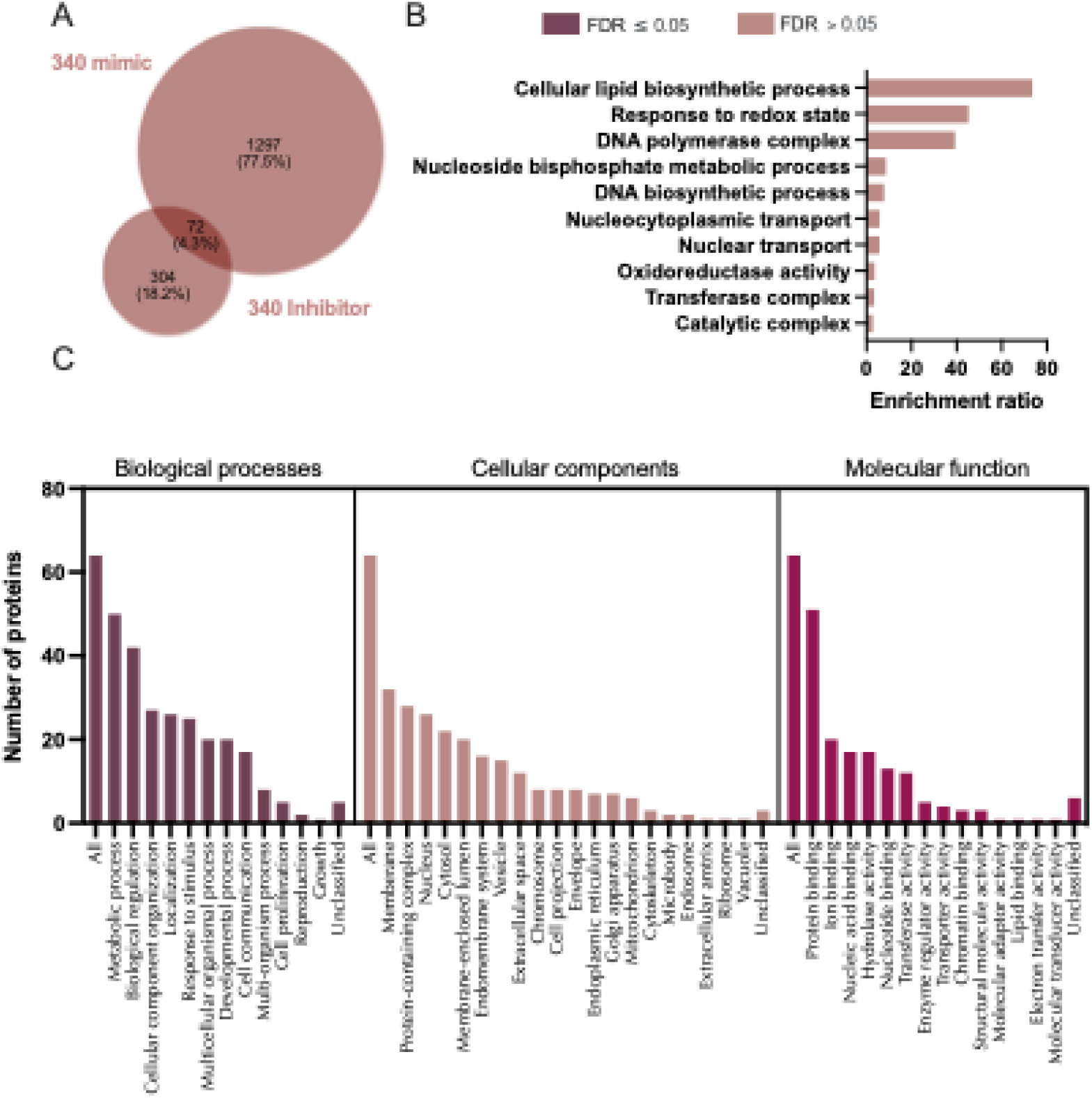
Proteins altered in Ishikawa cells following treatment with miR-340-5p mimic and inhibitor. **A)** Venn diagram depicting total number of significantly differentially expressed proteins (p<0.05) following transfection of Ishikawa cells (n=3 biological replicates) with miR-340-5p mimic (RHS) and inhibitor (LHS). B) Enriched KEGG pathways associated with miR-340-5p mimic and inhibition regulated proteins. C) WebGestalt overrepresentation analysis of biological process, cellular component and molecular function categories for identified significantly differentially expressed proteins (p<0.05) in response to miR-340-5p mimic and inhibition (total of 72).

We then sought to determine how many of these *in vitro* identified targets of mir-340-5p were predicted in mirDB. A total of 280 proteins were identified between predicted (mirDB) and confirmed (*in vitro*) targets (Supplementary Figure 3). Only 6.9% of total predicted targets were changed in endometrial cells treated with mir-340-5p mimic while 1.3% were altered following inhibition of mir-340-5p (Supplementary Figure 3: Supplementary Table 24). These proteins were overrepresented in pathways involving the nuclear pore, mRNA-3’-UTR binding, and cadherin binding (Supplementary Figure 3). Finally, we compared our *in vitro* mimic/inhibitor list of proteins, mirDB predicted targets, and those mRNAs altered by P4 in endometrial epithelial cells. In total, 171 proteins predicted to be targets by mirDB were altered *in vitro* by treatment with miR-340-5p mimic or inhibitor and were also altered by treatment of endometrial epithelial cells with P4. These proteins were involved in biological processes involving developmental processes, cell proliferation, and reproduction amongst others and are likely key components of P4 regulation of the endometrial epithelium (Figure 10, Supplementary Table 14).

**Figure 10.**
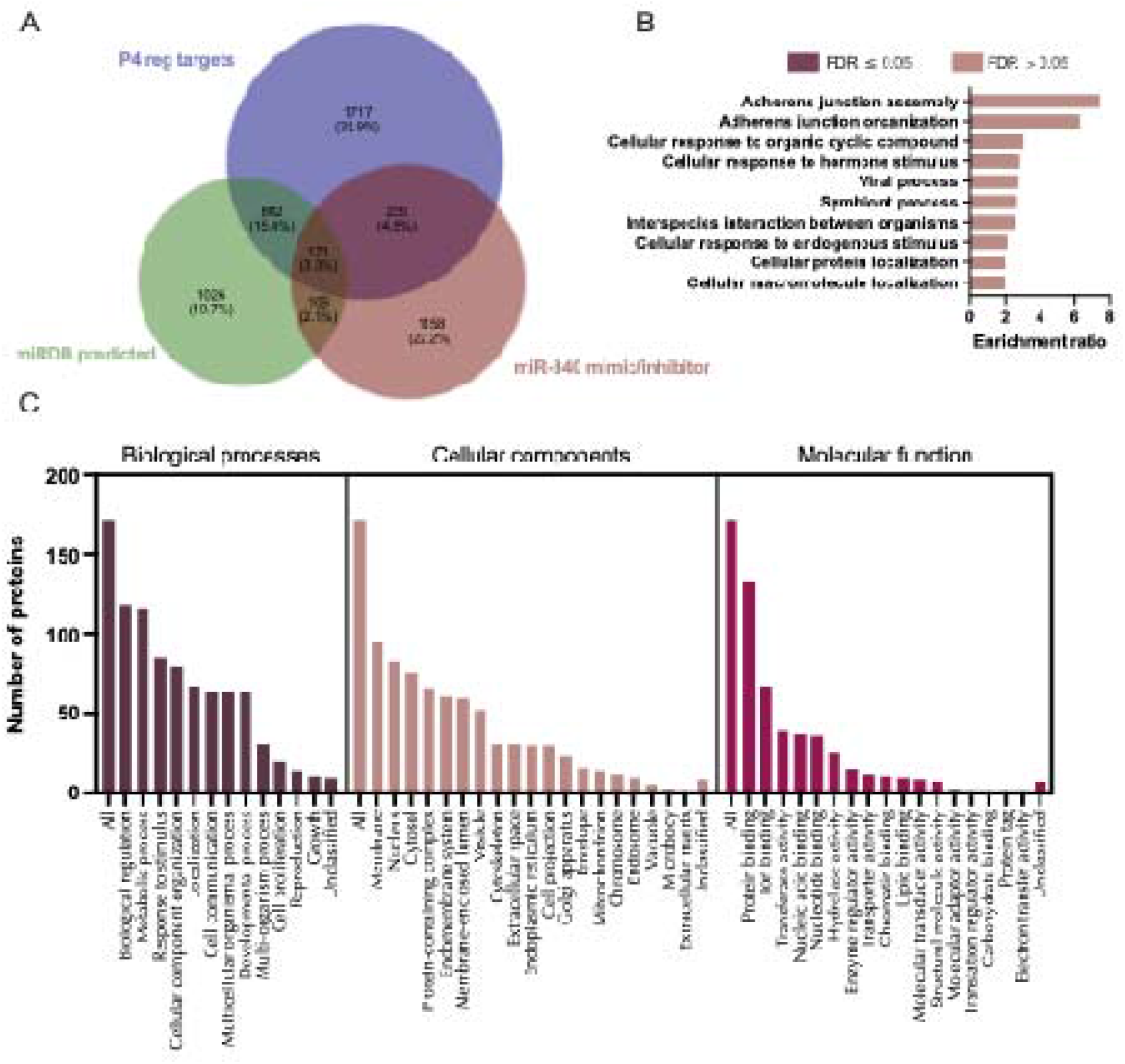
Proteins altered in Ishikawa cells following treatment with miR-340-5p mimic and inhibitor compared to mirDB predicted targets and P4 regulated mRNAs. **A)** Venn diagram depicting total number of significantly differentially expressed proteins (p<0.05) following transfection of Ishikawa cells (n=3 biological replicates) with miR-340-5p mimic or inhibitor vs non-targeting controls and mirDB predicted targets, or P4-regulated transcripts. **B)** Enriched KEGG pathways associated with miR-340-5p mimic and inhibition regulated proteins, P4-regulated mRNAs and predicted target overlap. C) WebGestalt overrepresentation analysis of biological process, cellular component and molecular function categories for identified significantly differentially expressed proteins (p<0.05) in response to miR-340-5p mimic and inhibition, predicted targets, and P4-regulated mRNAs overlap.

We wished to investigate whether the mRNA counterparts of proteins changed in response to miRNA mimics and inhibitors were present in RNASeq data from human endometrial biopsy samples (30). Of the proteins altered by miR-340-5p mimic, 94% of their corresponding mRNAs were detected in patient samples. Furthermore, only 13 of 1251 mRNAs were absent from 1 or more of n=36 samples. Ninety percent of proteins affected by inhibition of miR-340-5p were identified in RNASeq data, with only 5 of 326 mRNAs not present in all of the 36 patient samples. There were 59 of each of these sets of proteins which were changed by both mimic and inhibitor whose mRNA was found in patient samples (Figure 11, Supplementary Tables 15A-D).

**Figure 11:**
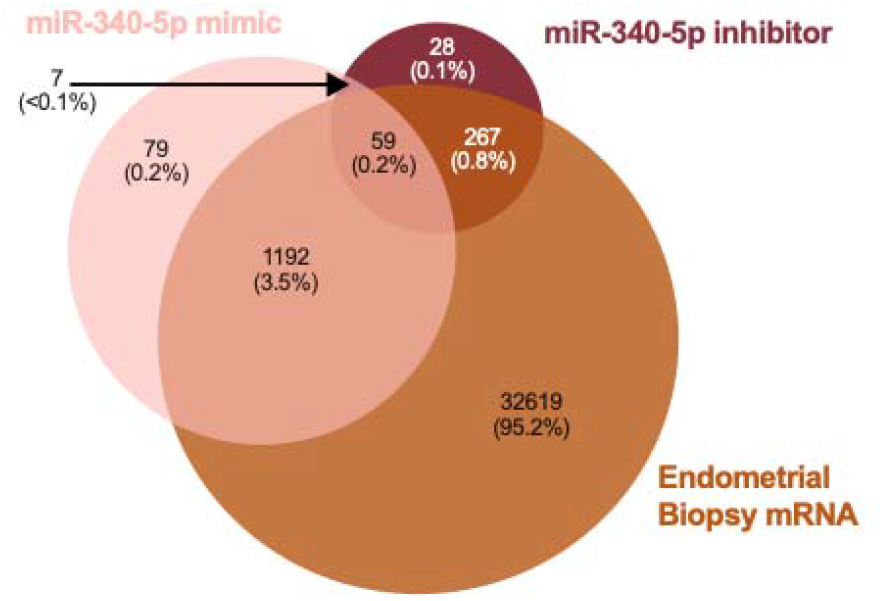
Comparison of proteins altered in abundance following transfection of Ishikawa cells (n=3 biological replicates) with miR-340-5p mimic or inhibitor compared to RNASeq data from human endometrial biopsies (n=36 biological replicates).

### In vitro targets of miR-542-3p in endometrial epithelial cells

Exclusion of off target effects identified 1,378 proteins altered by treatment of endometrial epithelial cells with miR-542-3p mimic (Supplementary Figure 4; Supplementary Table 25). These targets were overrepresented in pathways involved in ribonucleoprotein complex biogenesis and various metabolic processes (Supplementary Figure 4). Inhibition of miR-542-3p altered 975 proteins including those implicated in pathways related to RNA splicing, mRNA processing, and cellular protein localisation (Supplementary Figure 5, Supplementary Table 26). A core of 200 proteins were differentially abundant in response to treatment with both miR-542-3p mimic and inhibitor treatment (Figure 12, Supplementary table 16). These proteins were enriched pathways involving organelle and golgi inheritance as well as protein folding (Figure 12).

**Figure 12.**
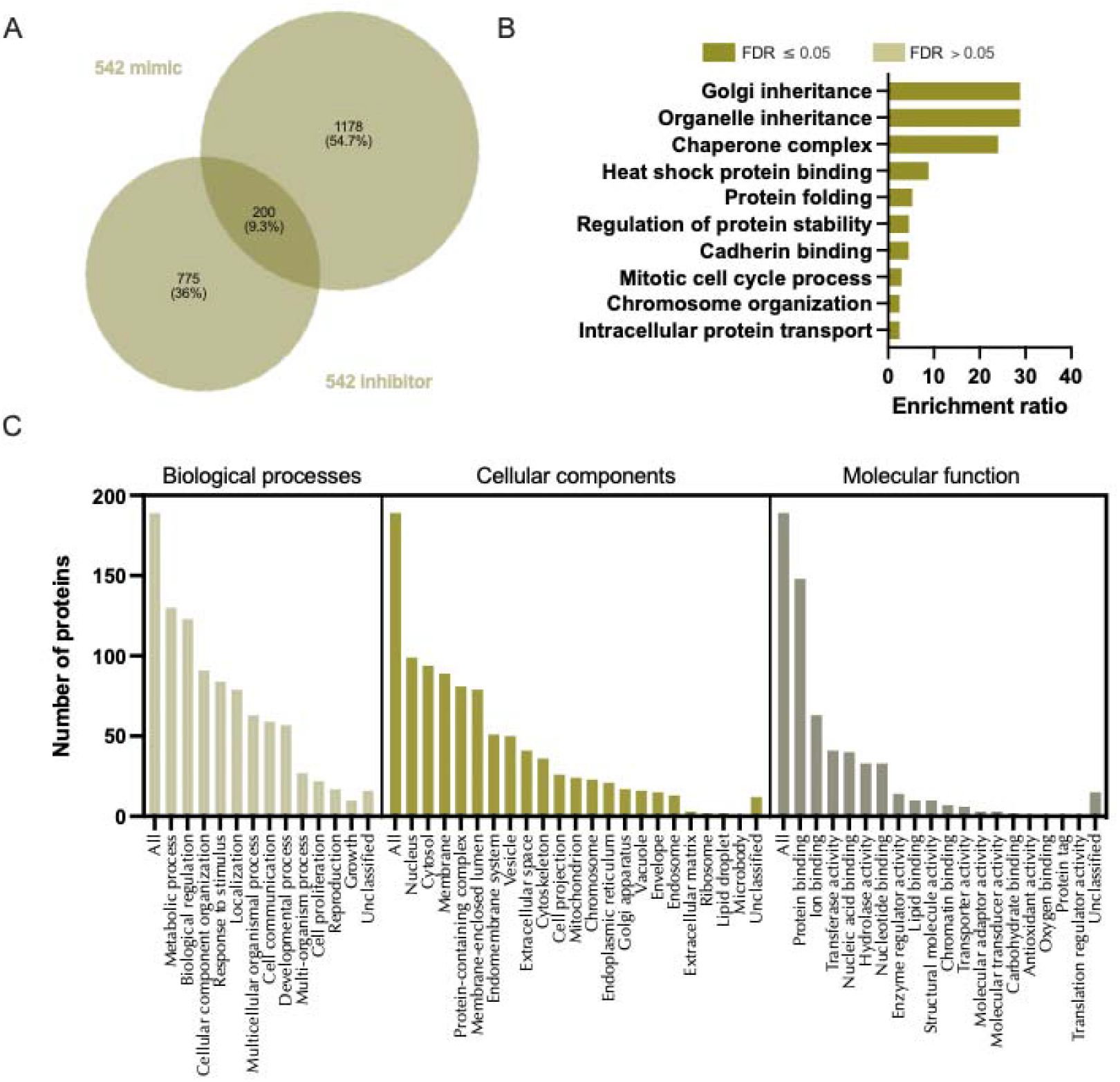
Proteins altered in Ishikawa cells following treatment with miR-542-3p mimic and inhibitor. **A)** Venn diagram depicting total number of significantly differentially expressed proteins (p<0.05) following transfection of Ishikawa cells (n=3 biological replicates) with miR-542-3p mimic (RHS) and inhibitor (LHS). B) Enriched KEGG pathways associated with miR-542-3p mimic and inhibition regulated proteins. C) WebGestalt overrepresentation analysis of biological process, cellular component and molecular function categories for identified significantly differentially expressed proteins (p<0.05) in response to miR-542-3p mimic and inhibition (total of 200).

On comparing these *in vitro* identified targets of miR-542-3p to those predicted by mirDB we identified 100 protein targets in common (Supplementary Figure 6, Supplementary Table 27). These were then compared to P4-regulated mRNAs and were assessed for the following criteria: 1) identified significantly differentially expressed proteins (p<0.05) in response to miR-542-3p mimic and/or inhibitor, 2) mirDB predicted targets of miR-542-3p, and 3) progesterone regulated miRbase targets of miR-542-3p (Figure 13, Supplementary Table 17). In total, only 46 proteins fit satisfied all three of our criteria and there are no significantly enriched pathways associated with these 46 proteins, however they appear to be enriched in thermogenesis (Figure 13).

**Figure 13.**
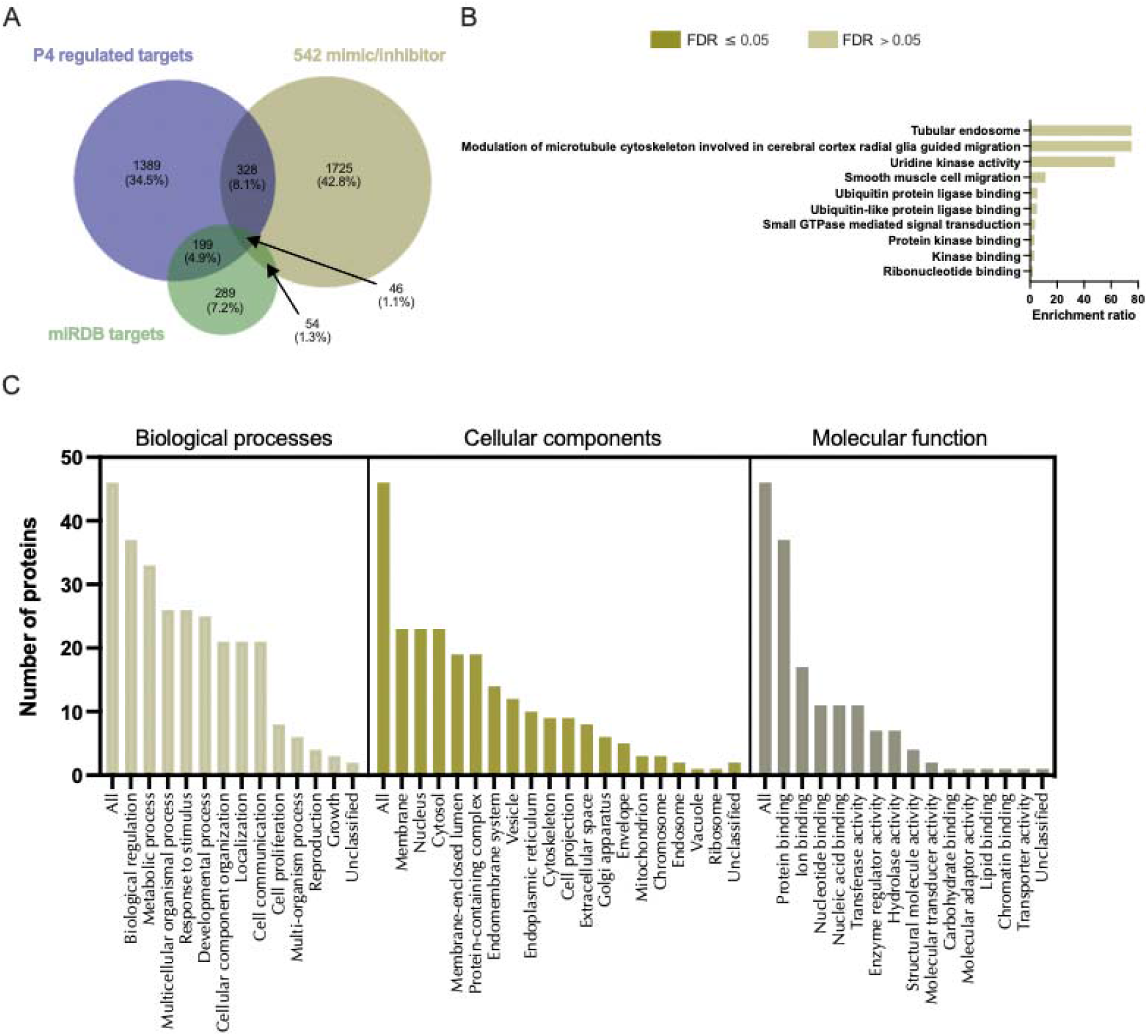
Proteins altered in Ishikawa cells following treatment with miR-542-3p mimic and inhibitor compared to mirDB predicted targets and P4 regulated mRNAs. **A)** Venn diagram depicting total number of significantly differentially expressed proteins (p<0.05) following transfection of Ishikawa cells (n=3 biological replicates) with miR-542-3p mimic or inhibitor vs non-targeting controls and mirDB predicted targets, or P4-regulated transcripts. **B)** Enriched KEGG pathways associated with miR-542-3p mimic and inhibition regulated proteins, P4-regulated mRNAs and predicted target overlap. C) WebGestalt overrepresentation analysis of biological process, cellular component and molecular function categories for identified significantly differentially expressed proteins (p<0.05) in response to miR-542-3p mimic and inhibition, predicted targets, and P4-regulated mRNAs overlap.

We carried out a comparison of proteomic data to human endometrial biopsies to determine whether proteins altered by miR-542-3p mimic and/or inhibitor were present in clinical samples (30). It was found that 94% of proteins changed in response to miR-542-3p mimic mRNA could be identified in RNASeq data. Of these 1261 proteins, only 11 were absent from 1 or more of n=36 samples. RNASeq data showed presence of mRNA corresponding to 95% of proteins altered in abundance by miR-542-3p inhibitor, with 8 of these 895 mRNAs not recognised in all 36 patients’ endometrial biopsies. A total of 181 proteins common to both mimic and inhibitor altered lists were found in RNASeq data (Figure 14, Supplementary Tables 18A-D)

**Figure 14:**
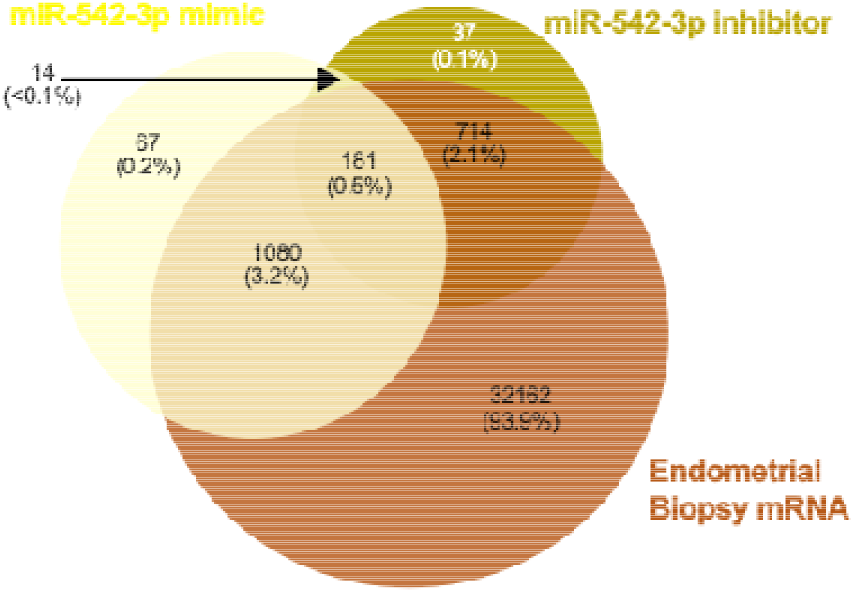
Comparison of proteins altered in abundance following transfection of Ishikawa cells (n=3 biological replicates) with miR-542-3p mimic or inhibitor compared to RNASeq data from human endometrial biopsies (n=36 biological replicates).

### In vitro targets of miR-671-5p in endometrial epithelial cells

Excluding off target effects, treatment of endometrial epithelial cells with a miR-671-5p mimic, 1,252 proteins were significantly changed (p<0.05) following exclusion of off target effects (Supplementary Figure 7; Supplementary Table 13). These proteins were involved in processes involving protein transport and localization (Supplementary Figure 7). Inhibition of miR-671-5p altered 492 proteins enriched in functions such as rRNA metabolic process, and generation of precursor metabolites and energy (Supplementary Figure 8; Supplementary Table 13). The number of proteins that were changed in response to both miR-671-5p mimic and inhibitor was 97, with overrepresented of functions related to response to endoplasmic reticulum stress (Figure 15; Supplementary Table 28).

**Figure 15.**
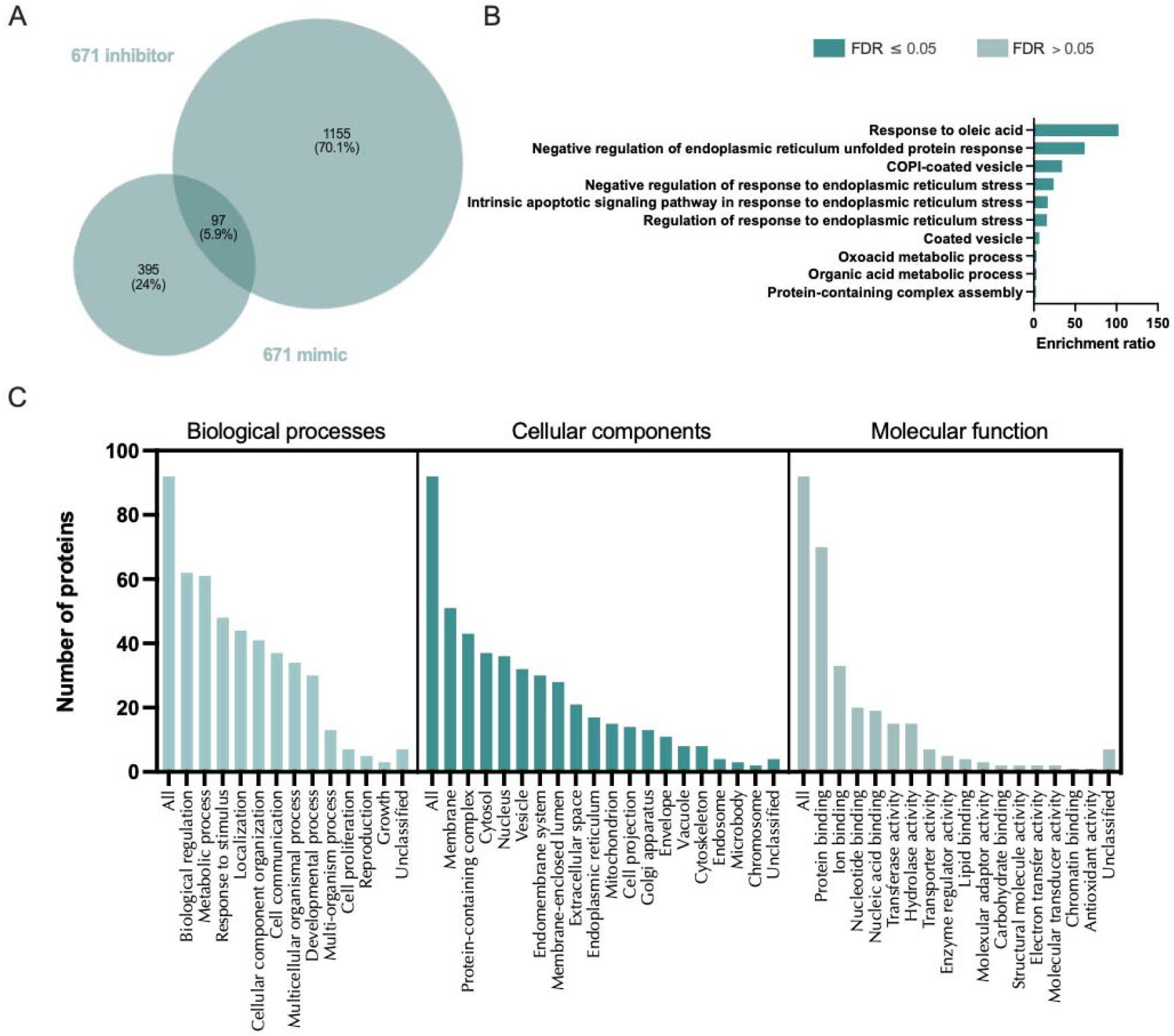
Proteins altered in Ishikawa cells following treatment with miR-671-5p mimic and inhibitor. **A)** Venn diagram depicting total number of significantly differentially expressed proteins (p<0.05) following transfection of Ishikawa cells (n=3 biological replicates) with miR-671-5p mimic (RHS) and inhibitor (LHS). B) Enriched KEGG pathways associated with miR-671-5p mimic and inhibition regulated proteins. C) WebGestalt overrepresentation analysis of biological process, cellular component and molecular function categories for identified significantly differentially expressed proteins (p<0.05) in response to miR-671-5p mimic and inhibition.

Comparing the set of proteins that were significantly changed following treatment with miR-671-5p mimic or inhibitor to those predicted to be targets of miR-671-5p, identified 95 proteins in common (Supplementary Figure 9; Supplementary Table 30). These proteins were significantly over-represented (FDR < 0.05) for processes involving the Golgi apparatus, cytosolic, intracellular, and vesicle-mediated transport (Supplementary Figure 9). Finally, by comparing proteins that fit each of the following 3 criteria: 1) identified significantly differentially expressed proteins (p < 0.05) in response to miR-671-5p mimic and/or inhibitor, 2) mirDB predicted targets of miR-671-5p, and 3) progesterone regulated miRbase targets of miR-671-5p discovers 46 that are common to all (Figure 16, Supplementary Table 20). The only significantly enriched category is in their cellular component organelle subcompartment (FDR < 0.05). Highly represented processes comprise various transport functions including golgi to lysosome (Figure 16).

**Figure 16.**
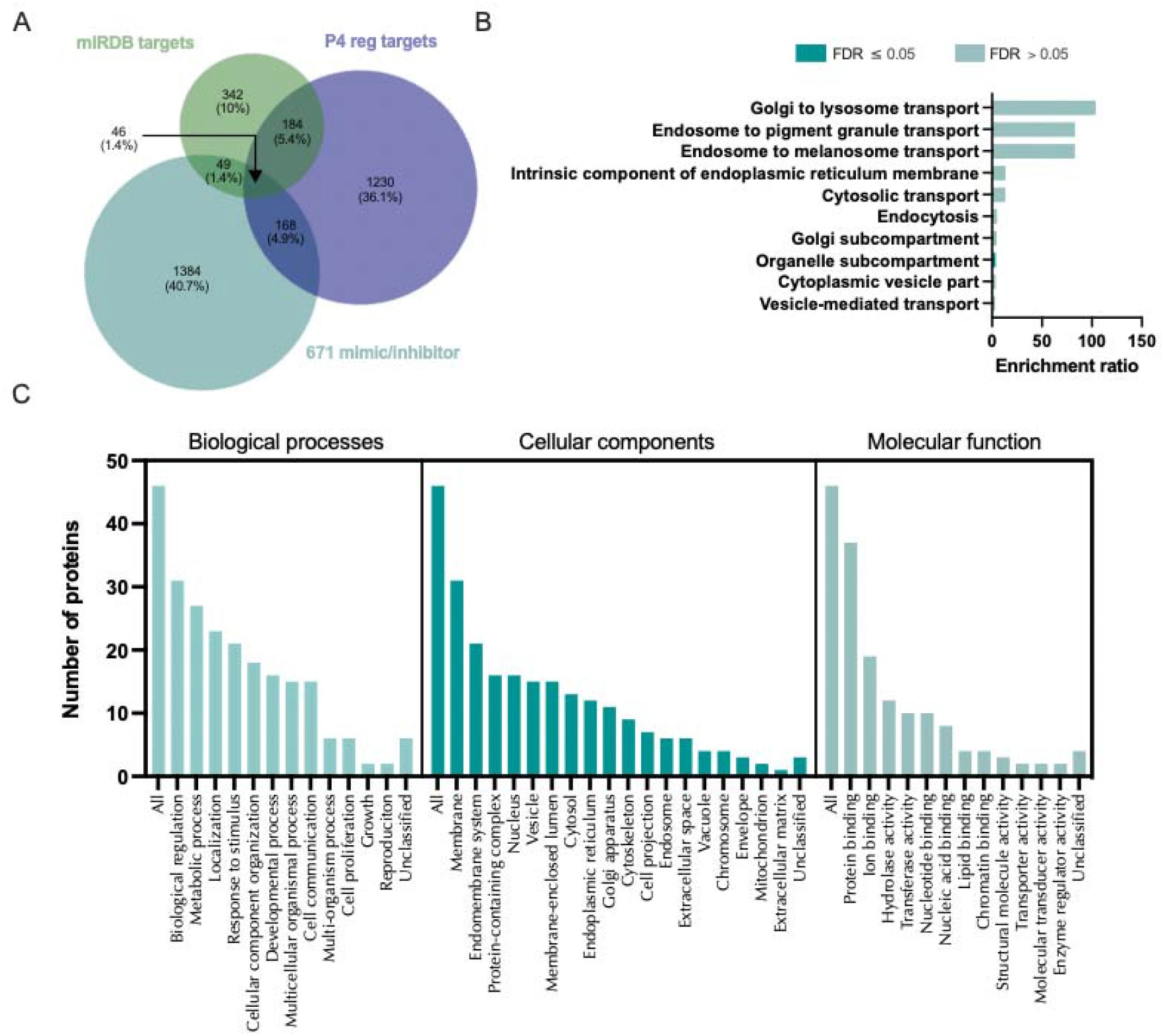
Proteins altered in Ishikawa cells following treatment with miR-671-5p mimic and inhibitor compared to mirDB predicted targets and P4 regulated mRNAs. **A)** Venn diagram depicting total number of significantly differentially expressed proteins (p<0.05) following transfection of Ishikawa cells (n=3 biological replicates) with miR-671-5p mimic or inhibitor vs non-targeting controls and mirDB predicted targets, or P4-regulated transcripts. **B)** Enriched KEGG pathways associated with miR-671-5p mimic and inhibition regulated proteins, P4-regulated mRNAs and predicted target overlap. C) WebGestalt overrepresentation analysis of biological process, cellular component and molecular function categories for identified significantly differentially expressed proteins (p<0.05) in response to miR-671-5p mimic and inhibition, predicted targets, and P4-regulated mRNAs overlap.

We examined presence of mRNAs in endometrial biopsies (30) to find those that correspond to proteins changed in abundance by miR-671-5p mimic and/or inhibitor. This resulted in identification of presence of mRNA for 94% of miR-671-5p mimic affected proteins. Of these 1150 mRNAs, only 8 were not found in all 36 endometrial samples. Ninety-five percent of proteins changed by the miR-671-5p inhibitor had corresponding mRNAs in patient samples and of those 449, only 8 were absent from 1 or more biopsy. Eighty-seven proteins were altered by both mimic and inhibitor as well as corresponding mRNA detected in endometrial biopsies (Figure 17; Supplementary Tables 21A-D).

**Figure 17:**
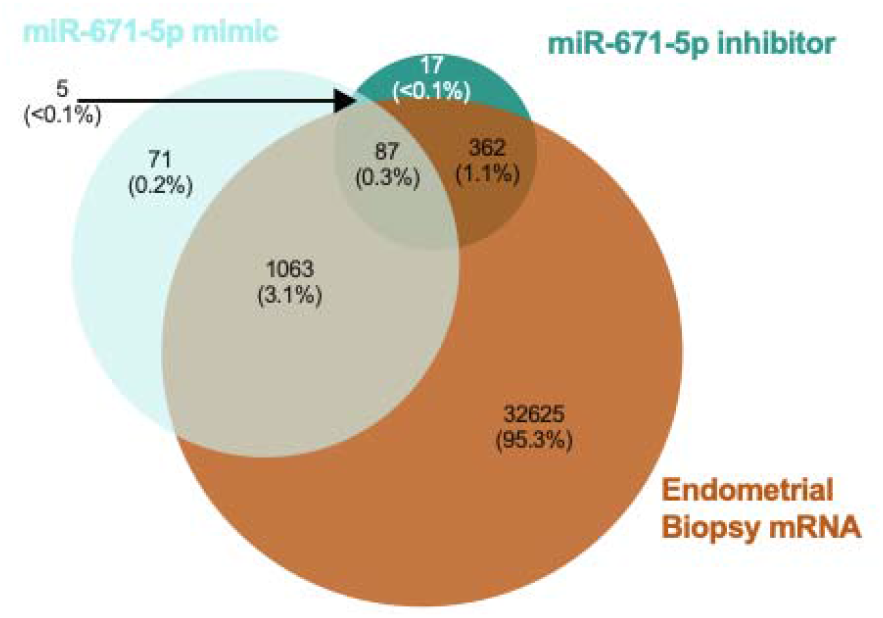
Comparison of proteins altered in abundance following transfection of Ishikawa cells (n=3 biological replicates) with miR-671-5p mimic or inhibitor compared to RNASeq data from human endometrial biopsies (n=36 biological replicates).

### Changes to miRNA profiles in patient samples with different clinical outcomes

Given the pathways these miRNAs regulate *in vitro we* sought to determine if their expression is dysregulated in endometrial biopsies from women who experienced pregnancy loss during the luteal phase of the cycle. We detected expression of all miRNAs in endometrial biopsies taken from patients during the luteal phase of their cycle, irrespective of prior or future pregnancy outcomes (Figure 18). No significant difference between patient cohorts was determined for 19 of the 20 mature seeds from the 13 microRNA families. However, expression of mir-340-5p showed an overall increase in expression in patients who had previously suffered a miscarriage and had a subsequent miscarriage, as compared to those who had infertility or previous miscarriage and subsequently went on to have a life birth outcome.

**Figure 18.**
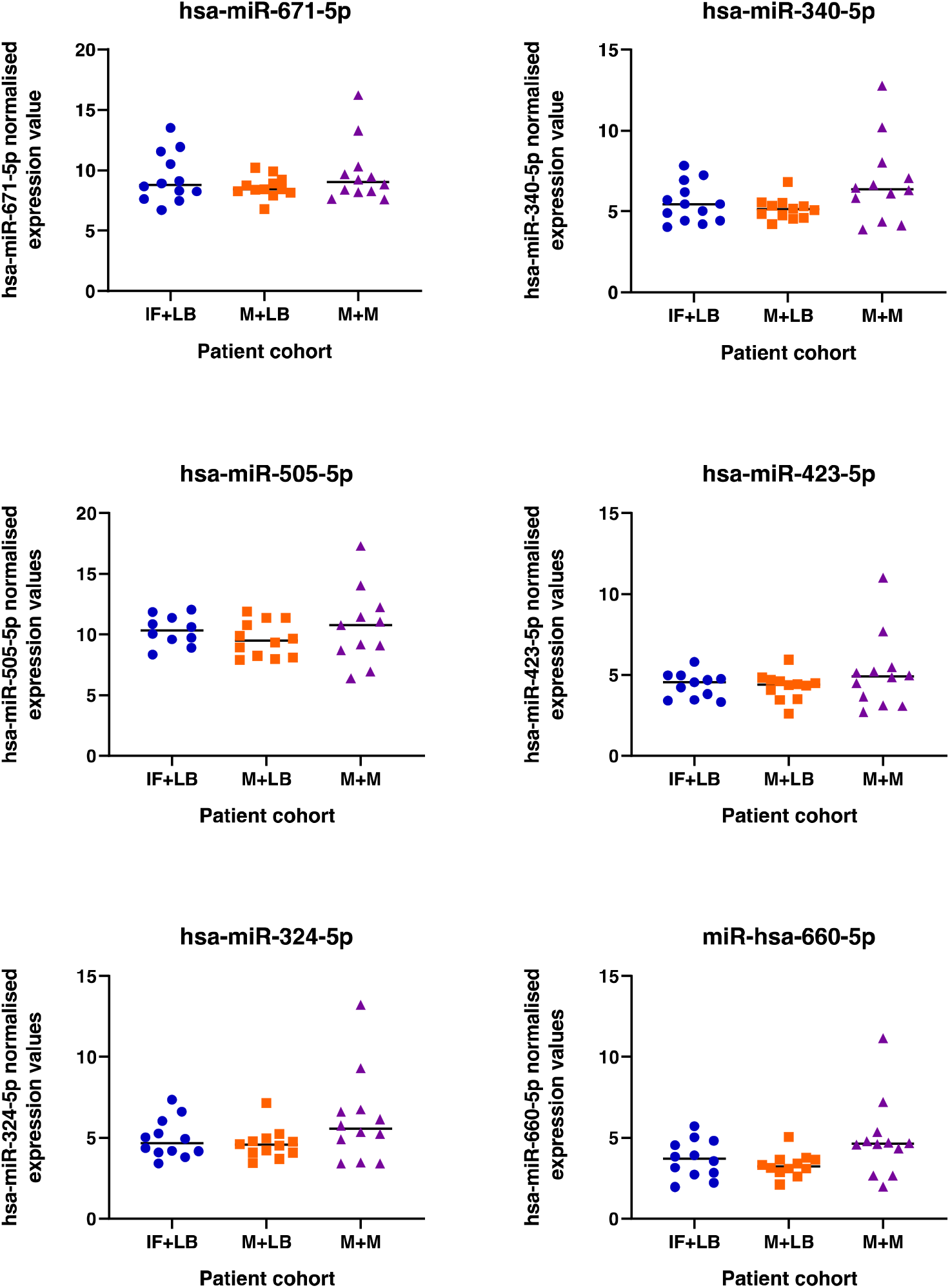
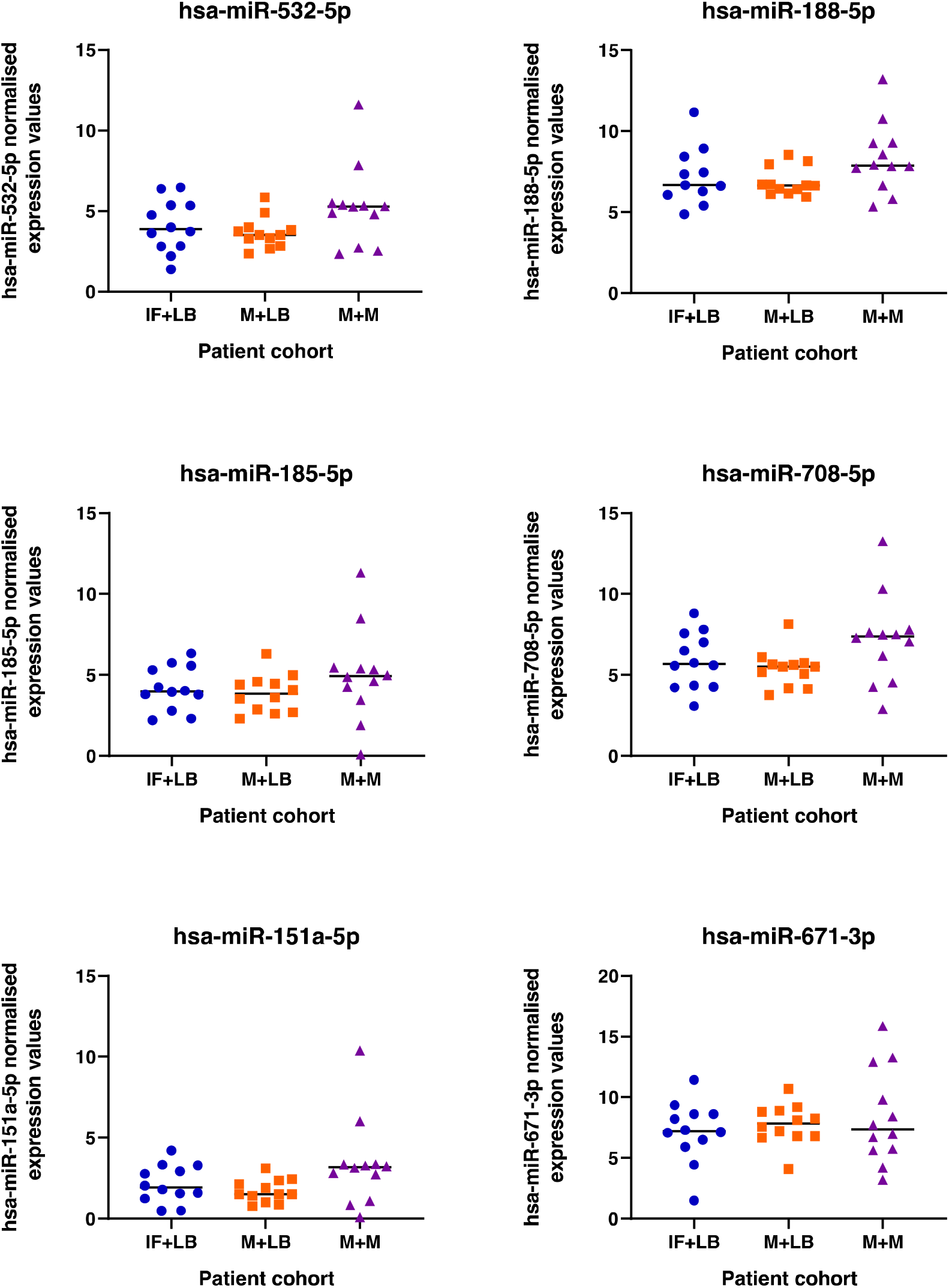
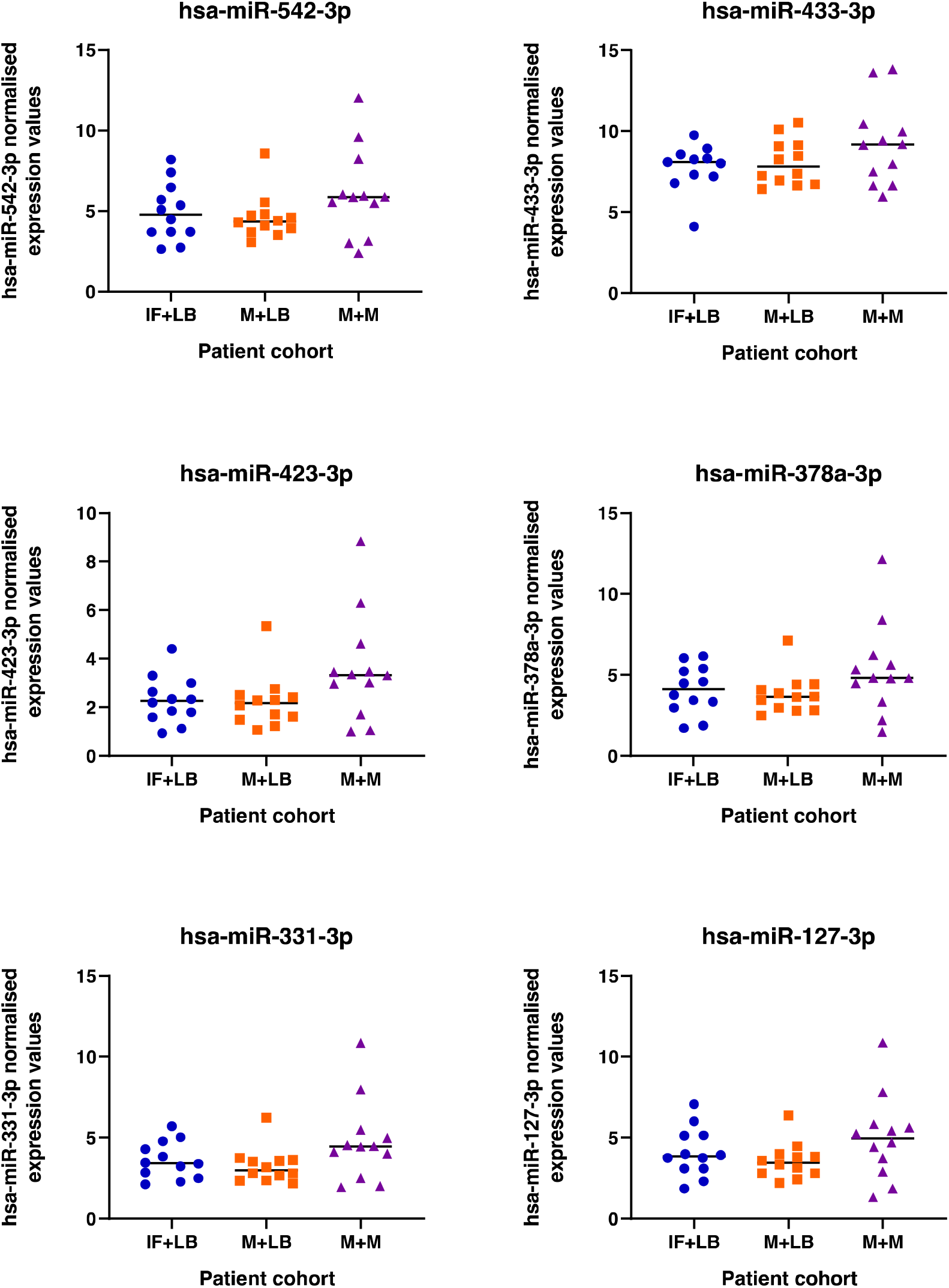
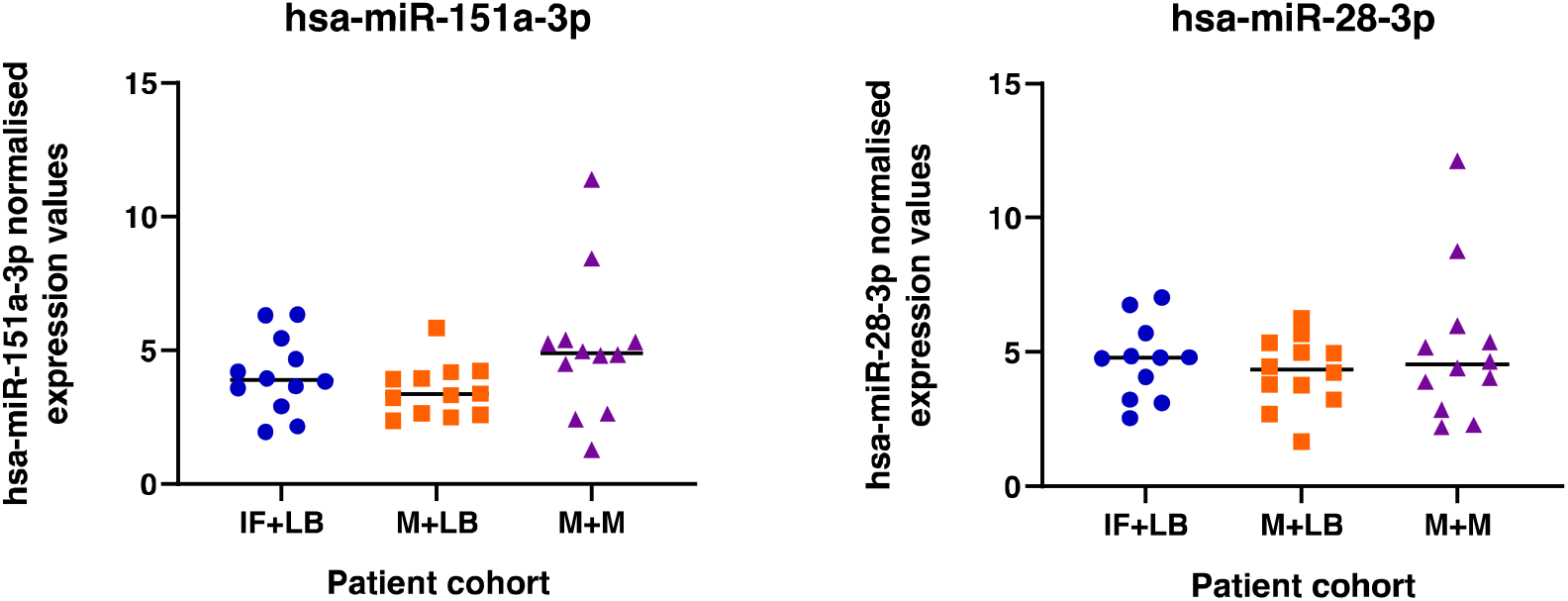
Expression of miRNAs in endometrial biopsy samples collected from patients during the mid-luteal phase of the menstrual cycle (LH+6-LH+9) from patients (n=12 per group) who have infertility and a subsequent live birth (IF+LB: blue dots), miscarriage and a subsequent live birth (M+LB: orange square), or miscarriage and a subsequent miscarriage (M+M: purple triangles). There was an overall effect on mir-340-5p expression (P<0.05) between groups.

## DISCUSSION

We tested the hypothesis that a set of miRNAs whose evolutionary appearance coincides with the emergence of placental mammals and were retained in all placental mammal lineages thereafter may play a role in uterine function for successful pregnancy in humans (18). The rationale for this is the fact that the endometrium and trophoblast have had to co-evolve regulatory signalling networks and the dysregulation of these networks may contribute to pregnancy loss. We have determined that miR-340-5p, miR-542-3p, and miR-671-5p were modified by exposure of endometrial epithelial cells to progesterone *in vitro*. These progesterone-regulated miRNAs occurred coordinate with alterations to selected mRNAs, a specific cohort of which they are predicted to target. Following treatment of endometrial epithelial cells with mimics or inhibitors for these three miRNAs we have determined the likely pathways that are P4 regulated by these miRNAs. We have also determined that the overwhelming majority of the proteins altered by these miRNAs are expressed in human endometrial biopsy samples (30). Moreover, we have demonstrated that there is a specific miRNA signature in the endometrium of a subset of women that is associated with miscarriage compared to those with a previous pregnancy loss but a subsequent successful pregnancy outcome. Collectively these data suggest these P4 modified and mammal-specific miRNAs, and the pathways in which their predicted targets function, are important for successful pregnancy and their dysregulation is associated with pregnancy outcome in a subset of women.

Recent studies have highlighted the transcriptional changes that occur in the endometrium during different stages of the menstrual cycle, but data on miRNA regulation of these processes – specifically in the endometrial epithelium - are more limited. miRNA profiling of whole endometrial biopsies (a heterogeneous mix of cell types) from the proliferative and mid-luteal phase of the menstrual cycle has revealed menstrual stagespecific miRNA signatures (31). This suggests the regulation of some of these 49 miRNAs is under the control of the steroid hormone environment at sampling. Similarly, our study has shown that P4 regulates expression of 3 of the 13 stem lineage miRNA gene families in endometrial epithelial cells suggesting these may contribute to regulation of the transcriptional response of the endometrium to changes in steroid hormones during the menstrual cycle.

The overall effect of P4 exposure to these epithelial cells altered transcripts involved in the processes of epithelium development and cellular proliferation, as well as functions involved in transcriptional regulation. Previous studies (31) found that predicted targets of miRNAs upregulated in mid-secretory phase endometrium were found to target cell-cycle regulators, possibly playing a role in cell turnover during the luteal phase. Analysis of the endometrium from women during the receptive stage of the menstrual cycle found ~ 21% of differentially expressed genes (653 out of 3,052) directly bind to or are regulated by the progesterone receptor (32). The functions related to these transcripts involve regulation of inflammatory response, estrogen response, cell death and interleukin/STAT signaling – some of which we identified as altered in expression following miRNA mimic/inhibitor treatment in our *in vitro* study. For example, addition of mimics for our three miRNAs of interest to Ishikawa cells changes expression of proteins involved in cellular proliferation (Fig 14).

Of the stem lineage miRNAs that were investigated we identified 3 that are modified by P4 *in vitro* in human endometrial epithelial cells, i.e., miR-340-5p, miR-542-3p, and miR-671-5p. We also showed via RNA sequencing that there were coordinate changes to some of the predicted targets of these miRNAs. P4 plays a critical role in endometrial function in a variety of placental mammals. Indeed, exposure of the endometrial epithelial and glandular cells to the sustained action of P4 is required to down-regulate its own receptor to facilitate receptivity to implantation – a mechanism that is conserved in all extant placental mammals studied to date (33). Analysis of the 473 mRNAs that were predicted targets of all three P4 modified miRNAs and were altered by P4 themselves (Figure 6) were involved in biological processes including cellular proliferation. In our *in vitro* treatment of endometrial epithelial cells with mimics/inhibitors of these P4 modified miRNAs, proteins involved in the biological processes of cell proliferation were also regulated by these miRNAs (Figures, 9, 12,15 panel C). Collectively we propose that P4 acting on endometrial epithelial cells, modifies expression of miR-340-5p, miR-542-3p, and miR-671-5p. This regulates expression of their targets that are involved in cellular proliferation to modify the endometrial epithelium to establish receptivity to implantation.

In addition to roles in cellular proliferation, these P4-regulated targets were also overrepresented in the molecular functions of ubiquitin-ubiquitin ligase activity and protein serine/threonine kinase activity. These are cellular processes (cell communication, proliferation, differentiation: (34)) that are also regulated by hCG treatment of human endometrial epithelial cells, as well as adhesion of these cells to molecules that make up the extracellular matrix (35). We have demonstrated these three miRNAs target proteins that function in adherens, and their assembly (miR-340), inter- and extracellular transport ((miR-340, miR-542, miR-671). This may indicate that P4 primes the endometrium for the further actions of hCG on the epithelium to facilitate secretion and transport of components of uterine luminal fluid and/or implantation – two key functions of the endometrial epithelium.

miRNAs have been shown previously to play a role in both successful pregnancy as well as pathophysiology. For example, miR-675 expression increases during placental development and inhibition of the expression of miR-675 negatively regulates cell proliferation in a mouse model of overgrowth (36). Several miRNAs play a role in trophoblast invasion (mir-16, −29b, −34a, −155, −210: (37)), are serum biomarkers of preeclampsia (38), and have been associated with recurrent pregnancy loss (mir-133a: (39)). Of the 13 stem lineage miRNA families investigated in this study, 5 (mir-154, mir-185, mir-188, mir-423, mir-542) have previously been implicated as differentially expressed in the placenta from preeclamptic pregnancies (40–42). Loss of mir-542 has also been implicated in endometrial stromal cell function by enhancing IGFBP1 expression in decidualisation (43) but this is the first report to examine expression in an epithelial model. We provide evidence of cell compartment-specific regulation of miRNA expression and their predicted targets – in keeping with the known physiological regulation of the endometrium [1]. To further substantiate this hypothesis, we used mimics and inhibitors to determine *in vitro* endometrial epithelial cell-specific targets of these P4-regulated miRNAs. The *in vitro* targets of these three miRNAs involved processes of transport and cellular localisation – key components of endometrial epithelial cell function. P4 stimulates the secretion of components of uterine luminal fluid (ULF) that supports growth and development of the conceptus prior to formation of the placenta. While there are limitations to the use of Ishikawa cells they do display some attributes of a receptive endometrial epithelial layer. More recently there have been developments in organoid models of endometrial glands and assembloids (44–46). Our approach, though a cellular monolayer of epithelium has allowed us to look at the potential mechanism by which these miRNAs may contribute to endometrial function *in vitro* and for the first time allow the comparison of computationally predicted targets with confirmed miRNA targets in this model. We propose that P4-regulated modification of these miRNAs and their targets function to provide components of the ULF that supports early pregnancy. Dysregulation of this process could contribute to RPL. In addition to identifying *in vitro* targets of these mimics, we compared these to computationally predicted targets of these miRNAs and found very limited overlap (Figures 10, 13, & 16). This is not surprising given the high false discovery rate known for target prediction software, of which mirDB is one of the better options (47).

We identified a difference in one of these P4-regulated stem-lineage miRNAs endometrial biopsies from women with different pregnancy outcomes. These samples are a heterogeneous combination of cells from endometrial biopsies. miRNAs have been associated with stromal cell decidualization in the human endometrium (48) and miRNA profiling of decidua from week 8 (elective termination) compared to those menstrual endometria - found 9 miRNAs to be differentially expressed including one stem lineage miRNA; mir-423 (49). This link between miRNAs and uterine receptivity was also established in a study of IVF patients who experienced recurrent implantation failure (RIF) (50) with dysregulation in the expression of a set of 13 miRNAs (not investigated in our study) in endometria of RIF compared to fertile women. Moreover, the protein targets of the 3 P4-modified miRNAs (miR-340, miR-542, and miR-671) were enriched for cell adhesion, cell cycle/ proliferation and inter- and -extracellular transport, all functions known to be implicated in embryo development/implantation (51). This further supports the important link between miRNAs and endometrial function. miRNA levels in blood plasma from women suffering from recurrent miscarriage to women who were not pregnant also differ including one of the stem linage miRNAs (miR-127-3p) (52). Although miR-127-3p is not altered by P4 in our study, there are many factors that could contribute to its modification in circulation, and it may be indicative of dysregulation of the endometrium at a different stage of pregnancy (52). In our study we have identified a subpopulation of endometrial samples that had aberrant miRNA expression profiles and this may identify a distinct cellular phenotype of endometrial from women who may be at risk of RM in the future. These could potentially provide targets for intervention in future.

In conclusion, we propose that 3 mammal-specific and evolutionarily conserved microRNA genes miR-340-5p, miR-542-3p, and miR-671-5p, along with some of their protein targets are regulated by progesterone – the key hormone of pregnancy in the endometrial epithelium in humans. Moreover, the regulation of these miRNAs and their protein targets regulate the function of transport and secretion, and adhesion of the endometrial epithelia required for successful implantation in humans. Dysfunction of these miRNAs (and therefore the targets they regulate) may contribute to endometrial-derived recurrent pregnancy loss in women.

## ACKNOWLEDGEMENTS

We would like to thank all of the women who provided endometrial biopsies for use in this study and the Biomedical Research Unit in Reproductive Health at UHCW for their support. This work was supported by N8 agri-food pump priming, QR GCRF, as well as BBSRC grant number BB/R017522/1 to NF as well as support from LTHT.

